# Wearable Neural Interfaces: Real-Time Identification of Motor Neuron Discharges in Dynamic Motor Tasks

**DOI:** 10.1101/2024.02.05.578874

**Authors:** Irene Mendez Guerra, Deren Y. Barsakcioglu, Dario Farina

## Abstract

**Objective:** Robustness to non-stationary conditions is essential to develop stable and accurate wearable neural interfaces.

**Approach:** We propose a novel adaptive electromyography (EMG) decomposition algorithm that builds on blind source separation methods by leveraging the Kullback-Liebler divergence and kurtosis of the signals as metrics for online learning. The proposed approach provides a theoretical framework to tune the adaptation hyperparameters and compensate for non-stationarities in the mixing matrix, such as due to dynamic contractions, and to identify the underlying motor neuron (MN) discharges. The adaptation is performed in real-time (∼22 ms of computational time per 100-ms batches).

**Main results:** The proposed adaptation algorithm significantly improved all decomposition performance metrics with respect to the absence of adaptation in a wide range of motion of the wrist (80°). The rate of agreement, sensitivity, and precision were ≥ 90% in ≥ 80% of the cases in both simulated and experimentally recorded data, according to a two-source validation approach.

**Significance:** The findings demonstrate the feasibility of accurately decoding MN discharges in real-time during dynamic contractions from wearable systems mounted at the wrist and forearm. Moreover, the study proposes an experimental validation method for EMG decomposition in dynamic tasks.

## 1 Introduction

The vision for neural interfaces is to tap the motor system to develop new control paradigms for rehabilitation, computer interaction, and augmentation by decoding user intentions from neural codes. One promising approach interfaces the pools of spinal motor neurons (MNs) – the last layer of the nervous system. By leveraging the electrical potential amplification of muscle fibres, MN activity can be decoded non-invasively from surface electromyography (EMG) using blind source separation [1, 2, 3] or deep learning techniques [4, 5]. In today’s rapidly evolving wearable landscape, the possibility to harness such MN activity for home rehabilitation or wide-range consumer electronics is closer than ever. Yet, a key remaining challenge for this vision, is the long-term stability of the EMG decomposition during non-stationary conditions.

While EMG decomposition algorithms have undergone rigorous validation, most approaches assume a mixing matrix that is constant over time. This simplification is valid during non-fatiguing isometric static contractions, where the joint position is fixed and the force is kept constant. In these highly controlled conditions, the firings of each active MNs are cyclostationary [6]. Each MN firing determines the generation of motor unit action potentials (MUAPs) at the neuromuscular junction. MUAPs then propagate along the membrane of the muscle fibres towards the tendon endings, producing a characteristic spatiotemporal electric potential. Decomposition algorithms seek to pseudo-invert these patterns to retrieve the underlying MN firings. To do so, blind source separation approaches first uncorrelate (whiten) the EMG signals and then iteratively find the separation vectors that maximise the sparsity of the MN spike trains [1, 2, 3]. Since the process is computationally intensive, decomposition parameters are usually calibrated in advance, prior to their real-time application in interactive settings [7]. Crucially, the effectiveness of such approach relies on the operational conditions remaining consistent with those present during the calibration phase. However, during natural movements, such stationary constraints are violated. When joints move, muscle fibres shorten and lengthen, changing not only the shape of the muscle but also pulling the surrounding tissue. These changes in the volume conductor alter the stereotypical profile of the MUAP waveforms, producing a dynamic mixing model.

At the wrist level, most forearm muscle fibres converge into tendons exhibiting unique electrical properties. When MUAPs reach the muscle-tendon interface, far-field potentials emerge as non-propagating components in the recorded signal before their complete extinction [8, 9, 10]. Similarly to the propagating components of the MUAPs, far-field potentials are also influenced by changes in fibre length and volume conductor properties [11].

Understanding the interplay between anatomical factors and dynamic modulation will contribute to the development of more robust EMG decomposition algorithms. Furthermore, these insights will enhance the design of wearable technologies, given that the wrist is the preferred location for users. Yet, only few studies have tried to characterised the modulation MUAPs undergo under dynamic contractions [12, 13, 14]. Most significantly, Kramberger et al. [13] evaluated MUAP changes from the biceps brachii across the entire elbow range of motion, showing a piece-wise linear trend with segments corresponding to 20% of the total elbow joint angle. These local linearities elucidate the reported ability to identify few MNs during dynamic contractions over relatively small ranges of motion under highly controlled experimental conditions [14, 15] and set the foundation for continuous updates.

Prior research has aimed to address the challenges of non-stationarities in the EMG mixing model by implementing recursive updates of the decomposition parameters based on new batches of data [16, 12, 17, 13, 18]. While these approaches have shown significant improvements in the accuracy of the detected spikes, the selection of some of the model hyperparameters is often ad-hoc based on the available dataset, providing a limited theoretical framework to deal with non-stationarities. In addition, validation so far has mainly relied on oversimplified simulated data [17] or hybrid data approaches [12, 13] due to the lack of better means. Only Yeung et al [18] used the gold-standard of matched intramuscular recordings for validations. However identifying common MNs in intramuscular and surface recordings remains challenging and limits the validation to few sources.

Recent advances in neuromuscular modelling offer new tools to address the validation problem. One notable example is NeuroMotion [19], which is a full-spectrum dynamic neuromechanical model that leverages physiological parameters from a myo-electric digital-twin [20], a muskuloskeletal model [21] and deep learning [22] to generate dynamic contractions with MN ground truth. While models like this one may provide previously unavailable insights on the modulation induced by dynamic contractions, it is still crucial to validate their outcomes experimentally.

Therefore, to develop a reliable and robust neural interface for long-term human-machine interaction, we first characterised how muscle fibre shortening and lengthening affect electric potentials using the NeuroMotion model [19] and experimentally recorded data from the forearm and the wrist. To analyse the experimental data, we propose a novel MUAP tracking approach for dynamic contractions that takes into account spatial, temporal, and amplitude features. Subsequently, we present an adaptive decomposition approach that recursively updates the decomposition parameters minimising the difference between the operational conditions and those during calibration. To do so, the algorithm leverages differentiable metrics such as the Kullback-Liebler divergence and kurtosis to guide the adaptation by minimising the median squared error with respect to the values at calibration. In this way, the proposed algorithm solely relies on signal properties providing a theoretical basis for the parameter optimisation. Here we show that the proposed adaptation can restore the accuracy of the decomposition under dynamic conditions in both simulated and experimental datasets after MUAP tracking in a double-source validation approach. Results show ≥ 90% RoA, sensitivity, and precision in ≥ 80% of the cases in both simulations and experimental data (median *>* 98% across all metrics and scenarios). Moreover, the execution time of the entire adaptation pipeline was lower than the average neuromechanical delay (∼ 200 ms [23]) making it suitable for real-time human-machine interaction applications.

## 2 Methods

### 2.1 Neuromuscular model simulations

Shortening and lengthening contractions at constant force were simulated using a full-spectrum dynamic neuromechanical model (NeuroMotion, [19]). The open-source model combines a neural model [24] that simulates MNs’ recruitment and force production; an AI-based volume conductor model (BioMime, [22]) that generates and/or morphs MUAPs based on dynamic physiological parameters; and an OpenSim upper limb musculoskeletal model to infer the dynamic parameters.

Briefly, the neuromuscular model simulates dynamic contractions by first emulating the desired movement in an upper limb musculoskeletal model [21]. The resulting muscle fibre length changes are fed into BioMime, an AI-based volume conductor model [22], along with the number of fibres innervated by each MN, their depth, angular position, innervation zone, and conduction velocity. The parameters related to muscles’ geometries and position are based on a myoelectric digital-twin [20] that generates accurate anatomical representations from magnetic resonance imaging. Notably, the model uses the changes in muscle fibre length to compute the corresponding changes in depth and conduction velocity under the assumption that muscles’ volumes are constant during dynamic contractions [25]. BioMime uses these parameters to generate the corresponding MUAPs and morph them across the simulated postural changes. Finally, EMG signals are generated by convolving the MN spike trains, simulated according to a MN recruitment and firing model [24], with their respective MUAPs at the corresponding posture. The obtained signals correspond to an electrode band arranged in 10 x 32 electrodes, that is placed around the proximal third of the forearm corresponding to the muscles’ bellies.

The simulated contractions involved an isometric index flexion while performing wrist flexion/extension to emulate the effect of postural changes of the hand. A pool of 100 MNs was simulated for the flexor digitorum superficialis muscle (FDS). The innervated fibres were randomly and uniformly distributed within the muscle volume, with innervation zones modelled as normally distributed around 50 ± 10 % (mean ± std) of the total fibre length. The MN’s recruitment thresholds and number of innervated muscle fibres were represented by an exponential function, which had a higher number of small low-threshold MNs compared to large high-threshold ones. Conduction velocities were drawn from a normal distribution (4.0 ± 0.5 m/s, mean ± std), and sorted according to the number of innervated fibres [26, 27]. When excitation surpassed a MN’s recruitment threshold, the MN began firing at a rate of 8 pps [24]. As the excitation increased, the discharge rate rose linearly by 3 pps for every 10% increase in excitation [24, 28]. The first MN recruited had its top discharge rate at 35 pps [24]. This rate reduced linearly with higher recruitment thresholds, up to 25 pps [24]. The last MN was recruited at 30% of the maximum excitation. Discharge rates followed a random Gaussian-distributed process, with a 0.2 coefficient of variation [24, 28].

To assess the effect of muscle shortening and lengthening, a FDS MUAP library was generated for each wrist joint angle degree (1 MUAP/°), ranging from -40°(flexion) to 40°(extension), corresponding to the range needed for most activities of daily living [29]. This process was repeated for 10 bootstrapping iterations, maintaining the same parameter distributions described above. Thereafter, two dynamic contractions of the wrist were simulated under 15% constant MN excitation. In them, the wrist joint angle followed a staircase pattern with constant phases every 10° from 0° to -40° for flexion or 40° for extension. The constant and transition phases lasted 30 s and 10 s, respectively. EMG signals were generated for each contraction at 2048 Hz sampling frequency and 30dB signal to noise ratio. Therefore, the simulated dataset comprised 20 contractions (10 bootstrapping sets, 2 conditions: flexion and extension). The constant phases of these patterns were used to calibrate a convolutive blind source separation (CBSS) [2] decomposition model (at 0°) and to evaluate the accuracy of the decomposition adaptation.

### 2.2 Experimental data

Nine healthy participants (four females, five males, ages: 24-30) took part in the study after signing informed consent forms. The experimental protocol was approved by the local ethics committee of Imperial College London (reference number 18IC4685) and was conducted in accordance with the Declaration of Helsinki.

EMG signals were concurrently recorded from the wrist (two electrode grids GR08MM1305, OT Bioelettronica, of 64 electrodes, arranged as 5 × 13, with 8 mm distance) and the forearm (three electrode grids GR10MM0808, OT Bioelettronica, of 64 channels arranged in 8 × 8 with 10 mm distance). First, the recording areas were shaved and treated with alcohol and abrasive gel. Thereafter, electrodes were placed along the circumference of the wrist right below the head of the ulna by visual inspection and physical palpation. Similarly, the electrodes at the forearm were placed around the circumference of the proximal third of the forearm corresponding to the muscles’ belly. EMG signals were acquired in monopolar mode (reference and ground bands were placed around the elbow) by a multi-channel amplifier (Quattrocento, OT Bioelettronica), bandpass filtered between 10 and 900 Hz, and sampled at 2048 Hz with 16-bit ADC precision. After acquisition, EMG signals were digitally bandpass filtered between 20 and 500 Hz (second-order Butterworth).

Participants were asked to place their right arm on a custom isokinetic system for wrist flexion and extension (see Figure 1). The isokinetic system included an elbow support, a motorised moving platform with hand support, and five adjustable micro load cells (0–5 kg CZL635, Phidget, 25 Hz sampling frequency) to record individual digit flexion and thumb abduction forces. While participants controlled the exerted finger forces, the angle of the hand platform was controlled by the motor (DOGA brushed geared, 24 V DC, 9 Nm, 45 rpm). The motor had integrated hall sensors that tracked the speed, direction and position of the motor, which were recorded at 100 Hz sampling frequency. To avoid any injuries, the motor operating range was limited with two inductive proximity sensors (LJ12A3-4-Z/BX, Heschen) between -50° and 50°, which was well below the maximum reported range of the wrist of ∼80° and ∼70° for flexion and extension, respectively [30]. Furthermore, during the experiment, an emergency stop button was held by the operator at all times. A custom Matlab (Mathworks Inc.) program was developed to synchronise all signal recordings and to provide participants with feedback of the cues and exerted finger forces, as well as the motor cue and current position.

**Figure 1:**
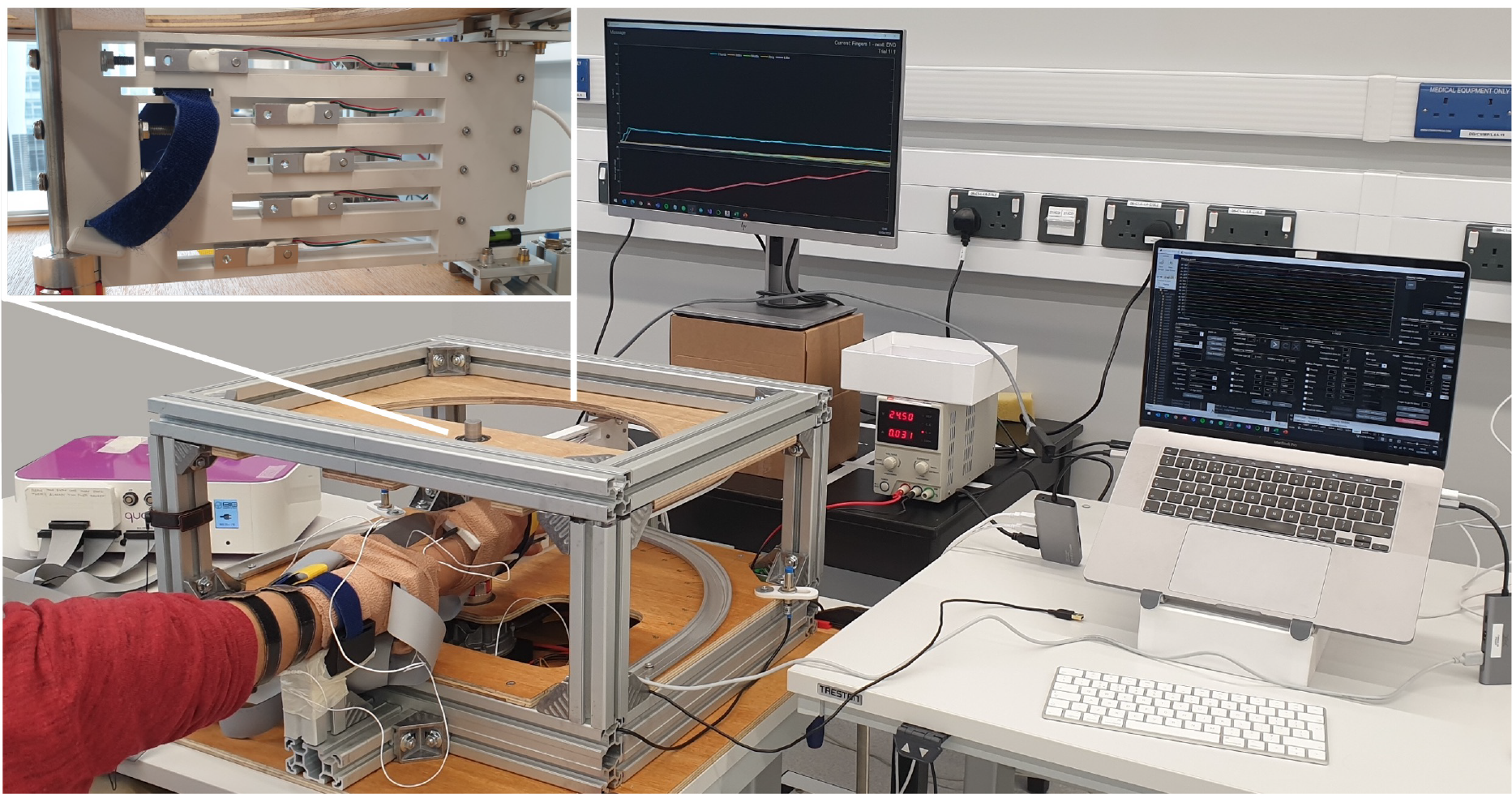
Experimental setup. EMG signals were concurrently recorded from the wrist and the forearm during isokinetic contractions where participants received feedback of their individual finger forces while a motorised platform controlled their wrist joint angle over flexion/extension. A zoomed in version of the motorised platform with the finger load cells is displayed on the top left corner.

At the beginning of the session, the hand was positioned so that the distal palmar crease was aligned with the rotating axis of the platform at the reference neutral position. At that point, the hand was securely strapped to the platform and the position of the force sensors was adjusted to ensure comfortable reach. Thereafter, the force sensor baselines and maximum voluntary contraction (MVC) forces were calibrated for each finger. Participants performed isometric digit flexions and thumb abduction at 15% MVC (individual fingers and all possible combinations of thumb, index, and middle) while the motor changed their wrist joint angle following a staircase pattern (from 0° to -40° flexion or 40° extension every 10° steps). To avoid participants’ fatigue, the constant and transition phases of the staircase angle pattern lasted 5 s and 10 s, respectively. Two repetitions of each contraction were recorded, resulting in 36 contractions in total (9 finger gestures, 2 conditions: flexion and extension, and 2 repetitions).

### 2.3 EMG generation model and decomposition

Briefly, EMG signals were generated by the spatiotemporal summation of the trains of potentials of the active MNs, which are mathematically formulated as the convolution between the MN spike trains and their corresponding MUAP waveforms [6]. EMG decomposition algorithms compute the pseudo-inverse of the MUAPs, under the assumption that MN spike trains are sparse (i.e. supergaussian distributions with positive kurtosis) and that each MUAP is unique for each MN, which has been also validated for the non-propagating far-field potential components at the wrist [31]. To simplify the formulation, the convolution between the volume conductor and the spike trains is unravelled into a linear matrix multiplication:

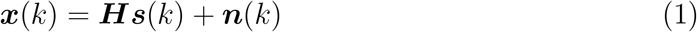

where ***x*** represents the extended (original and delayed versions) EMG signals; ***H***, the time unravelled MUAP waveforms; ***s***, the extended MN spike trains; ***n***, the extended noise; and *k*, the discrete time. In the absence of noise this results in the following inverse problem:

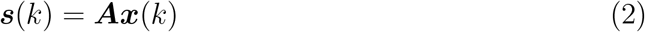

where ***A*** represents the pseudo-inverse of the extended MAUPs.

Thereafter, the extended EMG signals are centred by subtracting their mean and whitened so that their covariance is equal to the identity (i.e. are uncorrelated):

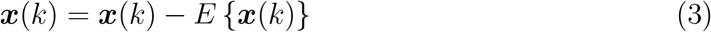

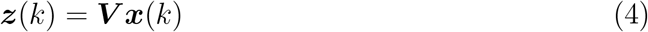

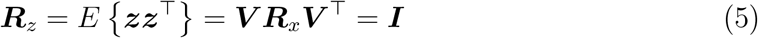

where *E*{·} is the expectation operator; ***z***, the whitened (extended and centred) EMG signals; ***V***, the whitening matrix; and ***R***_*z*_, the covariance matrix of the whitened signals. Whitening facilitates the identification of independent components by constraining the remaining transformation to an orthogonal space:

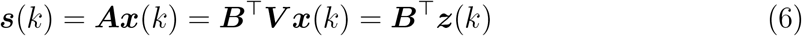

where ***B*** are the orthogonal separation vectors. These can be computed using fast fixed-point independent component analysis (fast-ICA) [32] for each source as previously proposed [2]:

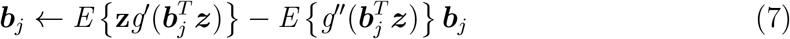

where ***b***_*j*_ represents the separation vector of the *j*^*th*^ updated source in each iteration; and *g*^*′*^(·) and *g*^*′′*^(·) are the first and second derivatives of contrast function *g*(·). Although fast-ICA maximises the statistical independence of the underlying sources, this assumption is violated by MNs due to the presence of common synaptic inputs within the same MN pool. In this case, instead, the maximised criteria is the sparsity of the MN spike trains, which is also optimised by common fast-ICA estimates of the negentropy (a measure of nongaussianity used as proxy for independence [32]) such the skewness, kurtosis, or their non-polynomial approximations [2]. To ensure the orthogonality of the resulting filters, Gram-Schmidt orthogonalisation is applied in each iteration:

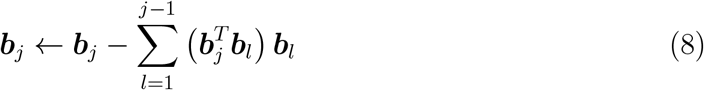

where the subscript *l* represents the previously updated separation vectors. In addition, the separation vector is normalised as:

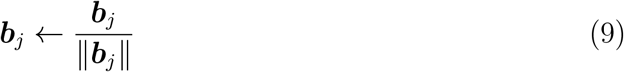

where ∥ · ∥ represents the vector norm operator. This iterative process is exited if the convergence tolerance or a maximum number of iterations is reached.

Finally, a second iterative process improves the quality of the detected MN firings based on the coefficient of variation of the interspike intervals. This involves K-means clustering for spike detection (baseline vs spike) and recalculating the separation vectors based on the detected spikes. In this case, the improvement process (often called reinforcement) ends once the coefficient of variation is minimised. Some approaches also include a reliability criteria to discard poorly decomposed MNs, such as the silhoutte measure (SIL) or pulse to noise ratio (PNR) that are surrogate indicators of the accuracy of the decomposition [33, 2].

### 2.4 Decomposition adaptation

In human-machine interfacing applications, EMG decomposition parameters are first calibrated from a sample contraction offline to ensure convergence. Thereafter, the whitening matrix, separation vectors, and spike detection parameters are applied in real-time on new testing data [7, 34]. This approach requires that the mixing matrices of both calibration and testing data remain identical and constant over time. While this assumption holds in isometric contractions, it is violated under postural changes or dynamic contractions. Hence, we present a new adaptation paradigm that updates the previously calibrated whitening matrix, separation vectors, and spike detection parameters by minimising the median squared error between the Kullback-Liebled divergence and kurtosis with respect to the calibration values for new batches of data (100 ms) to compensate for non-stationarities due to dynamic contractions or postural changes. The decomposition pipeline is shown in Figure 2.

**Figure 2:**
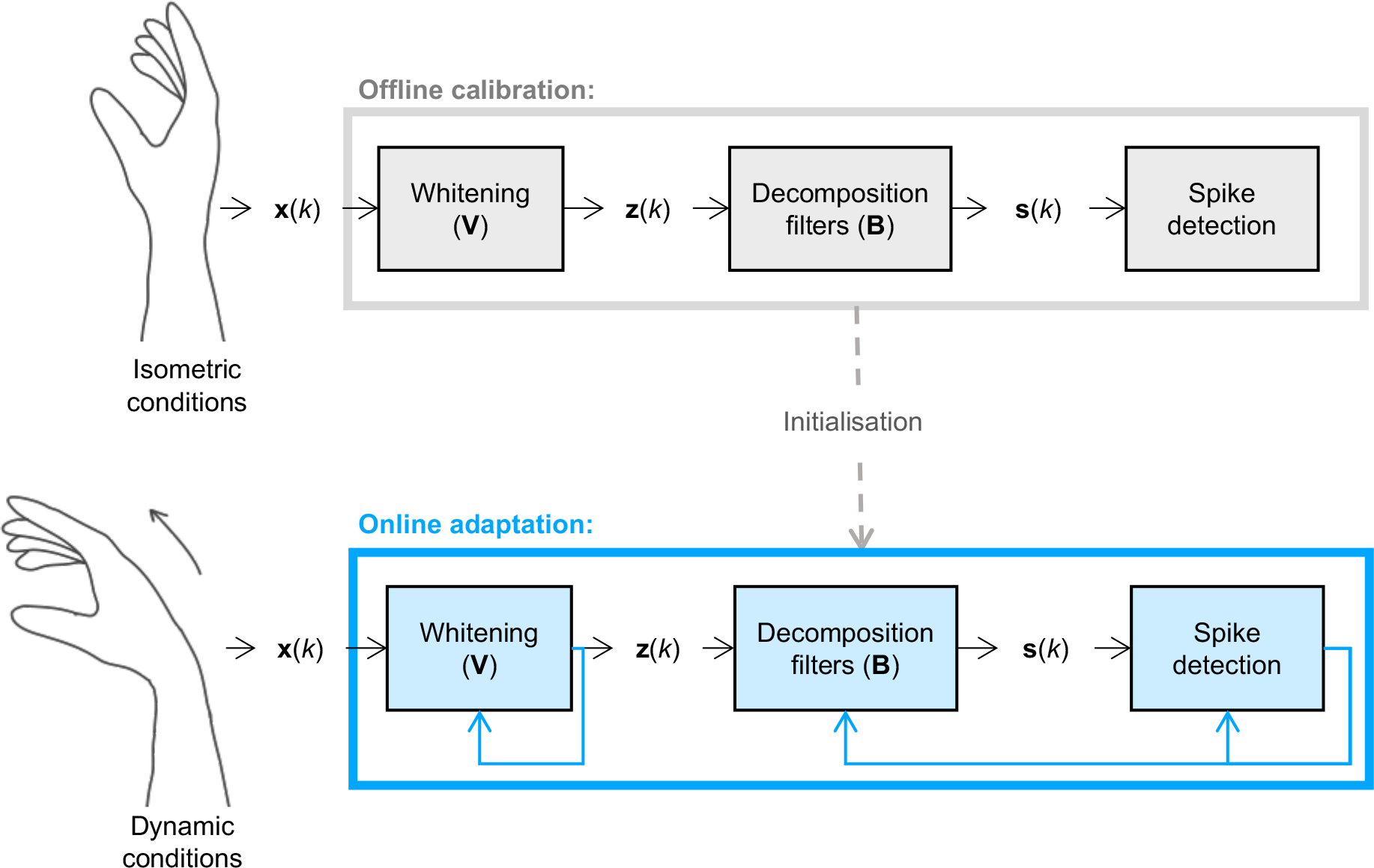
Proposed dynamic decomposition adaptation pipeline. A model previously calibrated during isometric conditions (top) is used to initialise the decomposition during dynamic contractions (bottom). In this case, the whitening, separation vectors, and spike detection parameters are updated in batches based on gradient descent.

#### 2.4.1 Whitening adaptation

The whitening matrix is often computed via eigenvalue decomposition [2] or singular valued decomposition [7]. However, these processes are not adaptive and computationally expensive to be run in real-time. To compensate for this, we propose a new adaptive approach based on the Kullback-Leibler divergence between two zero-mean normal distributions with covariance equal to ***R***_*z*_ and ***I*** [35]. The Kullback-Leibler is computed as:

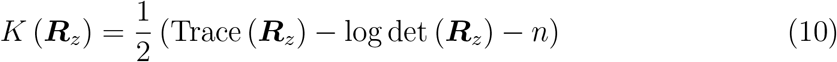

and serves as a measure of uncorrelation since it is equal to zero if ***R***_*z*_ = ***I*** and positive, otherwise. Therefore, the whitening matrix ***V*** is a minimiser of the following cost function *ϕ*_1_:

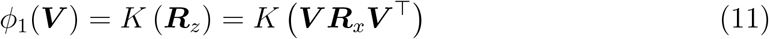

By taking the natural or relative gradient (see Appendix A in [35]) with respect to ***V*** we obtain:

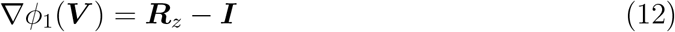

that results in the following whitening update rules:

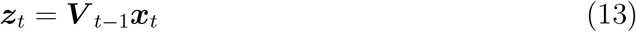

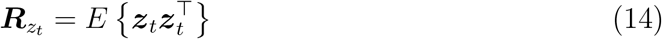

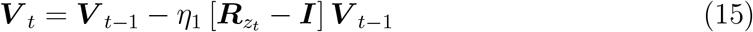

where *t* represents the batch number, and *η*_1_ is the learning rate that modulates the weight given to the new batch information. Its value is optimised as the minimiser of the median cost across all batches:

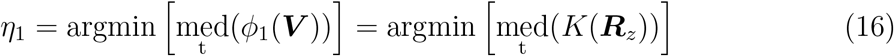

While it was found that this approach works well in simulations, for experimental data the minimum median squared error between the operational and calibration conditions had to be minimised due to differences in the topology of cost functions (Supplementary Figure 11), resulting in the updated cost function *ϕ*_2_:

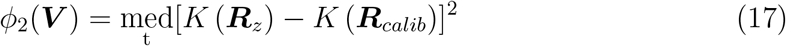

where ***R***_*calib*_ corresponds to the covariance of the whitened signals during the calibration phase. Similarly as in the previous case, taking the natural gradient results in the following update rules per batch:

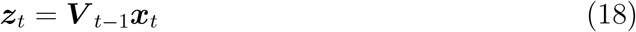

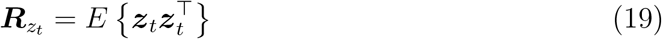

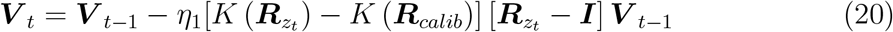

Since both the learning rate *η*_1_ and divergence error term are scalars, one can rewrite the previous Equation 20 as:

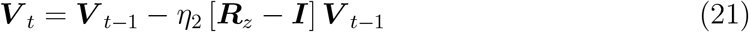

where *η*_2_ = *η*_1_[*K* (***R***_*z*_)−*K* (***R***_*calib*_)]. By assuming a constant error rate, Equations 15 and Equations 21 become analogous, and therefore the adaptation rule for both simulated and experimental data sets was the same, except for the choice of the hyperparameter (*η*_1_ for simulations, *η*_2_ for experimental conditions). In this case, *η*_2_ was optimised as the median squared error of Kullback-Liebler divergence across all batches with respect to the calibration:

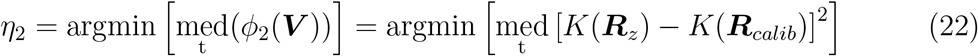

An initial grid-search optimisation resulted in the following hyperparameter values: *η*_1_ = 7*e*^−3^ for the all the simulated dataset, *η*_2_ = 1*e*^−3^ for the all the experimental wrist data, and *η*_2_ = 2*e*^−3^ for the all the experimental forearm data (see Supplementary Figure 11). After the optimal whitening hyperparameters were found, Equations 15 and Equations 21 were used to update the whitening matrix in simulations and experimental data, respectively.

#### 2.4.2 Adaptation of separation vectors adaptation

To adapt to nonstationarities in the separation vectors we propose the minimisation of the median squared error of the kurtosis of the sources with respect to their value at calibration. The kurtosis corresponds to the fourth-order cumulant of a random variable and is zero for gaussian distributions. Therefore, the kurtosis can be used as measure of nongaussianity, with higher values for more sparse sources ([36], pages 171-173). Indeed, its squared value is commonly used as approximation of the negentropy, although its nonpolynomial approximation is often preferred ([36], pages 183-185). Following this rationale, here we approximate the kurtosis (*κ*) per source (*s*) as:

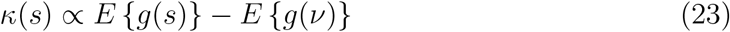

where *g*(*x*) = logcosh(*x*) represents a nonquadratic contrast function; and *ν* is a standardised gaussian variable. Given that *s* = ***b***^*⊤*^***z***, we can rewrite the previous expression in terms of the separation vectors resulting in:

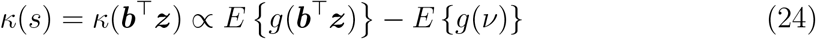

Therefore, the median squared error of the estimate of the kurtosis with respect to the calibration values can be expressed as:

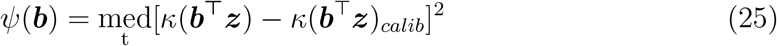

where *κ*(***b***^*⊤*^***z***)_*calib*_ represents the kurtosis of the sources detected during the calibration period. Taking the gradient relative to ***b***, we obtain the following update rule:

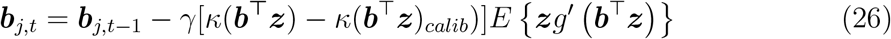

where *g*^*′*^(*x*) = tanh(*x*) is the first derivative of the contrast function *g*(*x*), and *γ* is the learning rate. By assuming a constant error rate as in the proposed whitening update, the previous equation can be rewritten into:

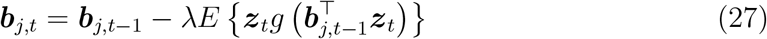

where *λ* = *γ*[*κ*(***b***^***⊤***^***z***) − *κ*(***b***^*⊤*^***z***)_*calib*_)]. Furthermore, since this update rule is applied separately to each source, we can also reduce the computational time by updating each separation vector at the times where a new spike is detected, leading to:

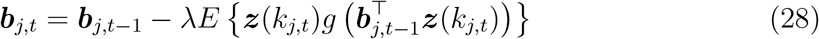

where *k*_*j,t*_ represents the newly detected spike times of the *j*^*th*^ source at the *t*^*th*^ batch. As in the proposed whitening adaptation, the hyperparameter (*λ*) was optimised as:

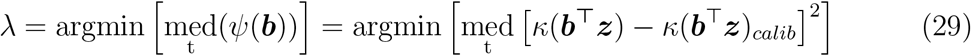

This optimisation resulted in *λ* = −3*e*^−3^ for the simulated dataset and *λ* = −5*e*^−4^ for both wrist and forearm experimental data. The change in sign of the hyperparameters makes Equation 28 equivalent to a maximisation of the estimated kurtosis regularised by the value of the initial conditions.

Importantly, after each batch update of Equation 28, the separation vectors were orthonormalised using Equations 8 and 9.

#### 2.4.3 Adaptation of spike detection

Since the separation vector approaches rely on the timings of the new spikes, an adaptive k-means algorithm, proposed in [17], was implemented for spike detection. This algorithm is briefly summarised by the following relation:

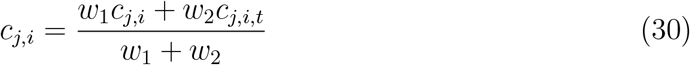

where *c*_*j,i*_ represent the k-means baseline (*i* = 0) and spikes (*i* = 1) centroids for the *j*^*th*^ MU; *c*_*j,i,t*_, are the newly computed centroids within the *t*^*th*^ batch exclusively; and *w*_1_ and *w*_2_ are weights empirically set to 5 and the number of newly detected spikes in the *t*^*th*^ batch, respectively. In this way, the adaptive k-means algorithm constitutes a weighted average of the previous and new centroids.

### 2.5 MUAP tracking

From the experimental data, we first labelled all MUAPs across angles with a tracking approach to reach a gold-standard for dynamics decomposition. For this purpose, first, e ach p lateau p hase o f t he s taircase w rist j oint a ngle w as i ndependently decomposed without adaptation using convolutive blind source separation [2]. Only reliable MNs with pulse to noise ratio (PNR) above or equal to 30 dB, discharge rate (DR) between 5-35 Hz (both included), and coefficient of va riation (C oV) of th e interspike intervals below or equal to 30 (%) were kept for each segment. The selected PNR threshold is equivalent to a *>*90% sensitivity and *<*2% false alarm rate in the decomposition accuracy [33]. Similarly, the accepted DR and CoV ranges relate to physiological values [24, 37, 38]. In addition, duplicated MNs within the same decomposition output (*RoA* ≥ 0.3) were also identified, and only the one with the highest PNR was k ept. The corresponding MUAPs were computed by spike trigger averaging on 25-ms windows. Thereafter, the MUAPs found at each angle were tracked to generate ground truth for the decomposition adaptation (Figure 3.a).

**Figure 3:**
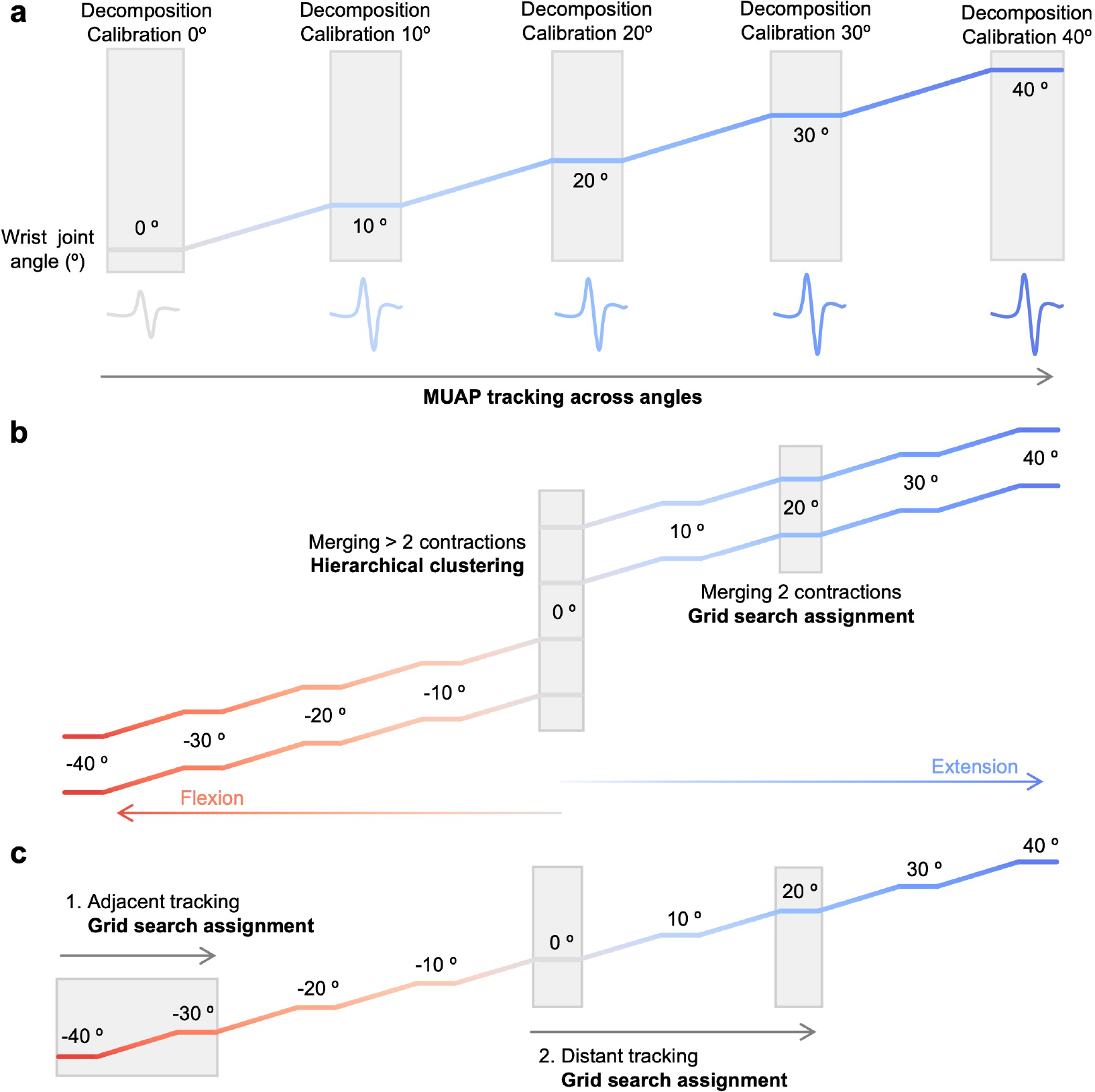
Experimental MUAP tracking approach. (a) Schematic of the motor unit action potentials (MUAPs) tracking approach after their computation at each wrist joint angle (colour coded) by independently calibrating a decomposition model at each plateau phase (shaded areas). (b) Merging of MUAPs across repetitions per angle (colour coded with flexion in red and extension in blue) based on their normalised mean squared error (NMSE) as metric for dissimilarity. Ward hierarchical clustering (new cluster formation at NMSE*>* 0.2) was applied when more than two contractions were being merged (at 0 °). When only two conditions were compared (rest of the angles), the proposed grid search assignment algorithm was applied instead (MUAPs were matched if NMSE*<* 0.2). The resulting library of unique MUAPs per angle was used for tracking across angles. (c) Tracking approach across angles using the proposed grid search assignment algorithm. (c.1) The approach sequentially compared adjacent angles to progressively track MUAP changes with the wrist joint angle (colour coded, same as in (b)). (c.2) A second loop compared distant angles to merge potentially disjoint tracking groups. In this case, only the MUAPs that had not been matched to their corresponding adjacent angle were included in the analysis. The matching threshold per comparison across angles was the same as in (b) for all cases.

When comparing MUAP pairs, these were first aligned by maximising their crosscorrelation over the union of channels with peak MUAP amplitude greater than:

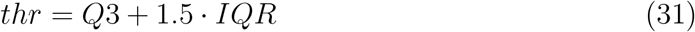

where *Q*3 represents the third quartile (75^*th*^ percentile), and *IQR* is the interquartile range (25^*th*^ to the 75^*th*^ percentile of the data). This selection effectively identifies the channels with outlying peak MUAP amplitude across the entire high-density electrode array, providing spatial selectivity to the comparison. Thereafter, the normalised mean squared error (NMSE) [39] was computed over the union of both sets of selected channels as:

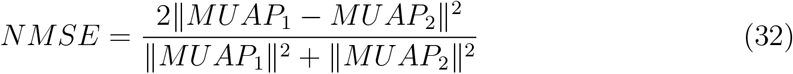

Note that NMSE is a dissimilarity metric with range [0, 2], where the equivalent similarity is expressed as 1 − NMSE. Importantly, the NMSE, unlike the correlation, can account for amplitude differences between the pairs of MUAPs, making it more suitable for dynamic contractions.

Thereafter a grid-search tracking algorithm was used to identify common MUAPs between two contractions. First, the NMSE matrix (*D*) with all pairwise comparisons between two MUAP sets of sizes *L* and *M* is computed. Then, the algorithm retrieves the set of values (*D*_*set*_) below a predefined NMSE threshold (*d*_*thr*_, which was set to *d*_*thr*_ = 0.2 in this study):

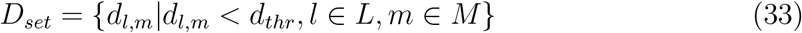

Every value in *D*_*set*_ is evaluated in ascending order, and a link between MUAPs *l* and *m* is formed if all the following conditions are met:

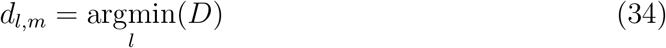

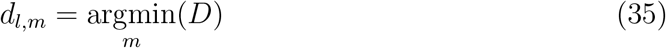

This approach was applied to merge MUAPs across repetitions for all angles except at 0 °, where four contractions were merged with Ward hierarchical clustering based on the NMSE with 0.2 threshold for cluster formation (Figure 3.b). When duplicated MUAPs were found, their spikes were inspected and the one with higher number of spikes, lower CoV and higher PNR was selected for further analyses. Once a library of unique MUAPs per angle was obtained, MUAPs were tracked across angles using the grid-search tracking algorithm sequentially. First, units were linked across consecutive angles; then the algorithm was applied across non-consecutive angles to merge disjoint groups of tracked MUAPs (Figure 3.c). In the disjoint angle comparison, only the MUAPs that had not been matched to their respective consecutive angle were included in the analysis. The corresponding group labels were propagated to all of the the previously identified duplicates across repetitions. In this manner, the sequential search approach enabled the effective tracking of MUAP modulations exceeding the NMSE threshold, and provided a validation set of spike trains for the experimental decomposition adaptation.

### 2.6 Decomposition adaptation evaluation

The decomposition adaptation pipeline was initialised with the whitening, separation vectors, and spike detection parameters computed from the first isometric phase of the simulated and experimentally recorded staircase contractions at 0°. Thereafter, the optimal adaptation hyperparameters were computed as described in Section 2.4. Once optimised, the decomposition adaptation was applied to the entire contraction. For the experimental recordings, only reliable and unique MNs (see section 2.5) were used to evaluate the decomposition adaptation. Similarly, for the simulated contractions, duplicated MNs within the same decomposition output (*RoA* ≥ 0.3) were also identified, and the one with the highest PNR was kept. Among those, only the decomposed MNs with *RoA* ≥ 0.9 with the simulated ones were used in further analyses.

To evaluate the accuracy of each adaptive pipeline, the rate of agreement, sensitivity, and precision were calculated as follows:

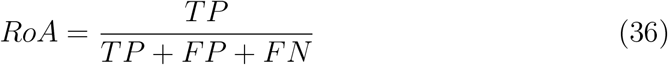

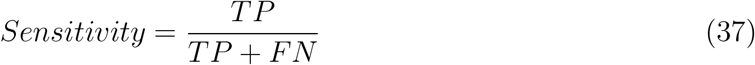

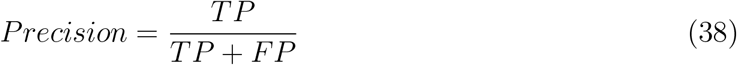

where *TP* represents the number of true positive spikes of a given MN (i.e. the spiked identified both in the adapted and in the tracked/simulated trains with a tolerance of ± 0.5 ms); *FP*, the false positives or the number of spikes detected in the adapted train only, and *FN*, the false negatives or the number of spikes detected in the tracked/simulated trains only. The PNR was also calculated to assess the effect of decomposition adaptation on the surrogate metric for decomposition accuracy in the absence of ground truth [33]. Finally, the NMSE was computed to evaluate the effect of the decomposition accuracy on the MUAP (dis)similarity.

### 2.7 Statistics

Statistical analyses were performed in SPSS 29.0 (IBM) with *p <* 0.05 level of significance. Nonparametric statistical test were used when the data was not normally distributed as assessed by the Shapiro-Wilk test (*p <* 0.05). The effect of adaptation in the decomposition performance metrics of the simulated dataset was assessed by the Wilcoxon signed rank test. Mann-Whitney U tests were used to evaluate differences in the number of detected MNs between locations (wrist vs forearm), as well as differences in their respective PNR, DR, and CoV. A generalised linear mixed model with repeated measures was used to evaluate the differences in NMSE across potentially matching MUAPs (matched vs second best alternative vs mean across all alternatives for a given angle pair; within-factor effect) and location (wrist vs forearm; between-factor effect). Similarly, individual generalised linear mixed models with repeated measures were used to analyse the effect of adaptation (within-factor effect) across locations (between-factor effect) for each decomposition performance metric in experimental conditions.

## 3 Results

### 3.1 Decomposition adaptation in simulations

Ten bootstrapping iterations of wrist flexion and extension from -40° to 40° were simulated using NeuroMotion. Figure 4.a shows snapshots of the muskuloskeletal model across the simulated wrist joint angles with the corresponding changes in muscle fibre length, depth, and conduction velocity (Figure 4.b-d, respectively). These non-stationary physiological parameters resulted in changes in the MUAP waveforms detected from the forearm with the wrist joint angle (Figure 4.e). Figure 4.f shows that these dynamics resulted in higher median NMSE during maximum flexion than extension (median (interquartile range): 0.17 (0.13-0.22) n.u vs. 0.09 (0.07-0.11) n.u., respectively). Importantly, these differences did not affect the identifiability of the simulated MNs as the NMSE across MUAPs of different MNs remained constant across angles (Figure 4.g).

**Figure 4:**
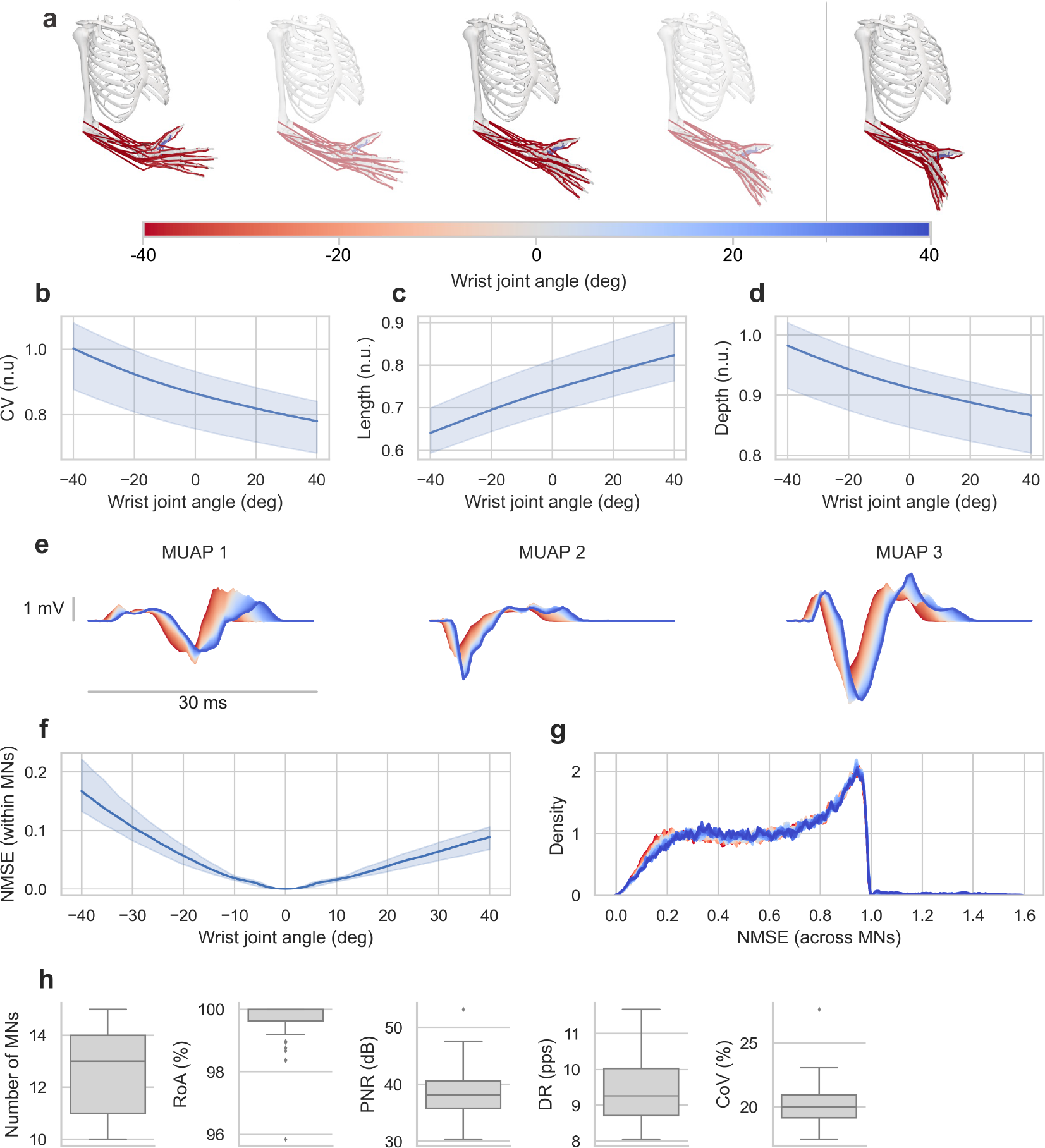
Dynamic neuromechanical simulations. (a) Snapshots of the muskuloskeletal model while performing a wrist flexion from -40°to neutral (0°) to extension up to 40°. Mean (solid line) and standard deviation (shaded area) of the induced modulations in muscle fibre length in normalised units (b) as informed by the muskuloskeletal model, and the corresponding changes in muscle fibre depth (c) and conduction velocity (CV) (d) in normalised units under the assumption of constant muscle volume during dynamics for all contractions. (e) One electrode visualisation of three generated motor unit action potentials (MUAPs) at the forearm with the above parameters for each wrist joint angle (colour). (f) Mean (solid line) and standard deviation (shaded area) of the normalised mean squared error (NMSE) within each motor neuron (MN) across wrist joint angles with respect to the neutral position (0°) for all the simulated MNs. (g) Distribution of the NMSE across MNs for each wrist joint angle (colours). (h) Number of identified MNs identified during the calibration phase per contraction, along with their rate of agreement (RoA) with the simulated ones, pulse to noise ratio (PNR),, discharge rate (DR), and coefficient of variation of the interspike intervals (CoV).

Out of the 100 simulated MNs per contraction, a median of 13 (11-14) MNs were identified during the decomposition calibration phase at 0° per contraction, with 100.00 (99.63-100.00) % median RoA with the simulated ones and 38.11 (35.85-40.59) dB median PNR, 9.26 (8.71-10.03) pps median DR, and 19.98 (19.15-20.91) % median CoV (Figure 4.h). Figure 5.a depicts the overall simulated wrist joint angle pattern for a representative wrist flexion contraction along with the MNs excitation level. The innervated pulse trains before and after adaptation for three MNs over the shaded region in Figure 5.a are depicted in Figure 5.b-d, along with the simulated and detected firings (top), and corresponding RoA across the entire contraction. Figure 5 illustrates the decrease in decomposition accuracy during dynamic contractions, where MN firings become undetectable after ∼10° without adaptation. The overall RoA, sensitivity, precision, NMSE, and PNR for all MNs across angles are displayed in Figure 6. In general, decomposition accuracy was completely lost after 20° in both flexion and extension when decomposition parameters were fixed. Notably, decomposition adaptation compensated this effect across all angles and metrics with significant improvements in the RoA (98.10 (94.42-99.16) % vs 35.86 (22.77-53.61) % for adaptation and no adaptation, respectively, *p <* 0.001), sensitivity (98.21 (95.25-99.20) % vs 36.46 (23.64-56.03) % for adaptation and no adaptation, respectively, *p <* 0.001), precision (99.92 (99.52-100.00) % vs 62.61 (48.17-80.00) % for adaptation and no adaptation, respectively, *p <* 0.001), NMSE (0.04 (0.03-0.06) n.u. vs 0.17 (0.06-0.32) n.u. for adaptation and no adaptation, respectively, *p <* 0.001), and PNR (39.08 (37.13-41.28) dB vs 30.68 (28.21-33.90) dB for adaptation and no adaptation, respectively, *p <* 0.001).

**Figure 5:**
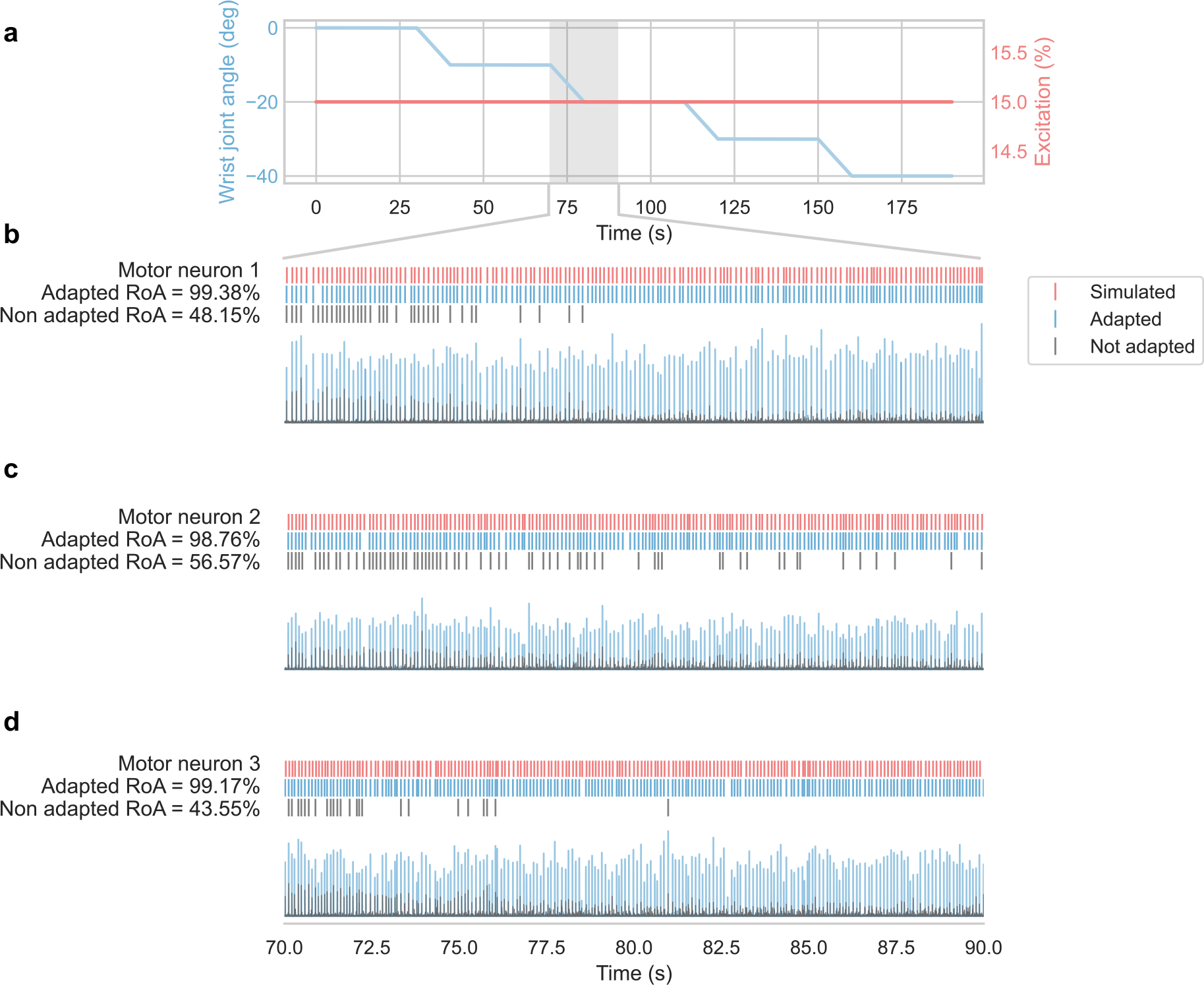
Detection of motor neurons (MNs) firings during simulated dynamics. (a) Simulated wrist joint flexion angle (blue) and MN excitation level (red) for a representative contraction. (b-d) detected MN firings before adaptation (grey) and after adaptation (blue) with the simulated ones for reference (red) for three representative MN over the shaded part of the contraction in (a). The respective rates of agreement (RoA) are shown on the left, and the innervation pulse trains are depicted below before adaptation (grey) and after adaptation (blue) for each unit. Panels b-d demonstrate the need for decomposition adaptation as MN firings become undetectable after ∼10° without adaptation.

**Figure 6:**
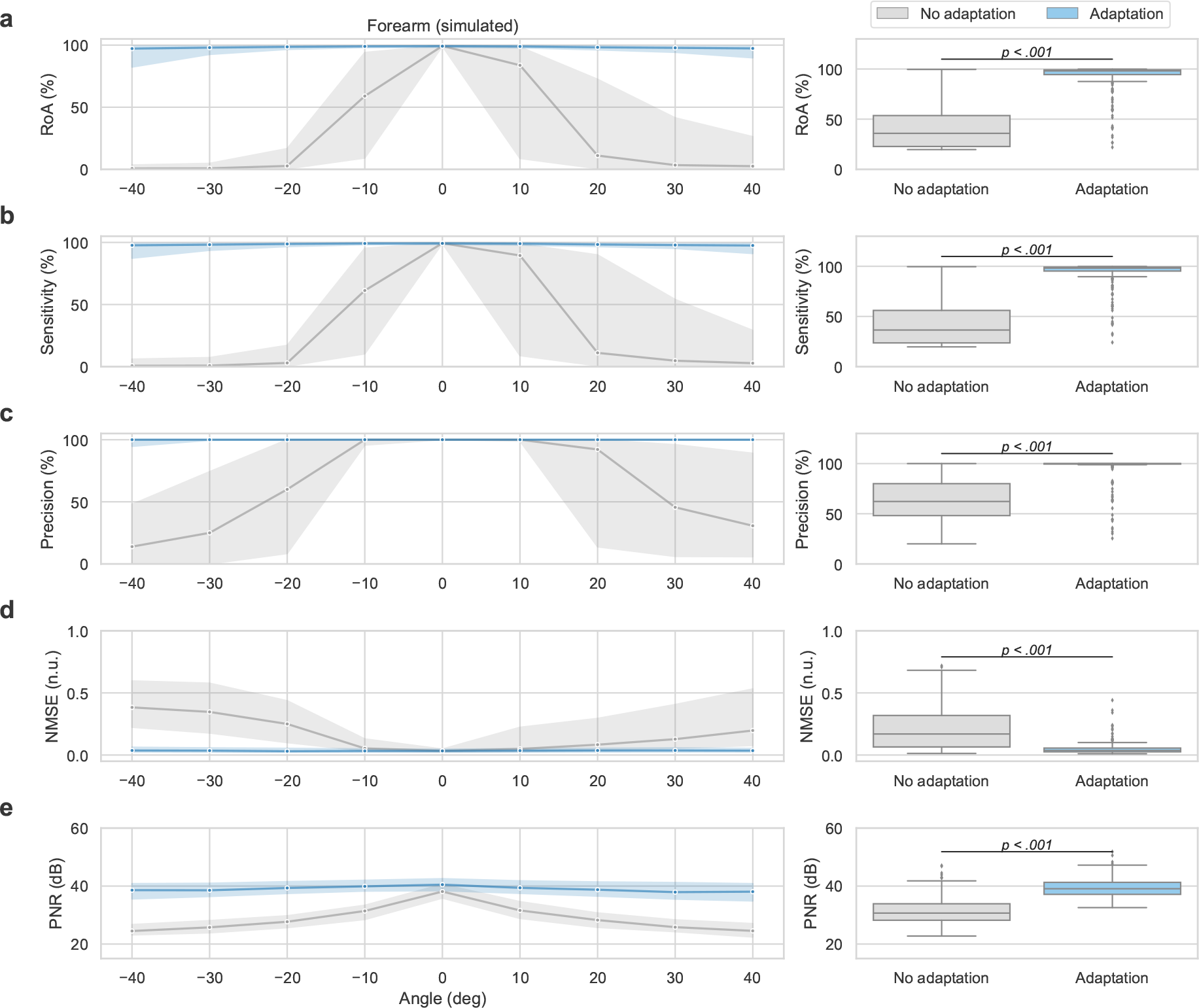
Decomposition adaptation performance in simulations. On the left, median (solid line) and interquartile range (shaded region) before (grey) and after (blue) decomposition adaptation across all simulated units for each angle for (a) rate of agreement (RoA), (b) sensitivity, (c) precision, (d) normalised mean squared error (NMSE), and (e) pulse to noise ratio (PNR). The right panels show the median and quartiles distribution of the metrics across all the simulated motor neurons (MNs). Note the different scale on the y-axis between left and right for visualisation purposes. Significant *p*-values are show in the plots as assessed by related-samples Wilcoxon signed rank tests. Decomposition adaptation significantly improved all performance metrics (*p <* 0.001).

The median execution time of the decomposition adaptation pipeline was 21.69 (20.65-22.60) ms per 100 ms batches for 320 electrodes (extension factor = 3) and 13 (11-14) MNs in a Intel Core i9, 2.4 GHz 8-Core, with 64 GB machine.

Taken together, these results show that the proposed decomposition adaptation algorithm could compensate the non-stationary changes induced by the simulated dynamic contractions in real-time.

### 3.2 Experimental MUAP tracking

Figure 7 shows two representative examples of the tracked MUAPs at the wrist and at the forearm with the grid-search algorithm, along with the tracking paths of MNs detected at different angles. In both cases, MUAP differences across angles can be appreciated in the non-propagating components of the wrist MUAPs, and around the innervation point of the forearm ones driven by the change in muscle morphology during dynamic contractions.

**Figure 7:**
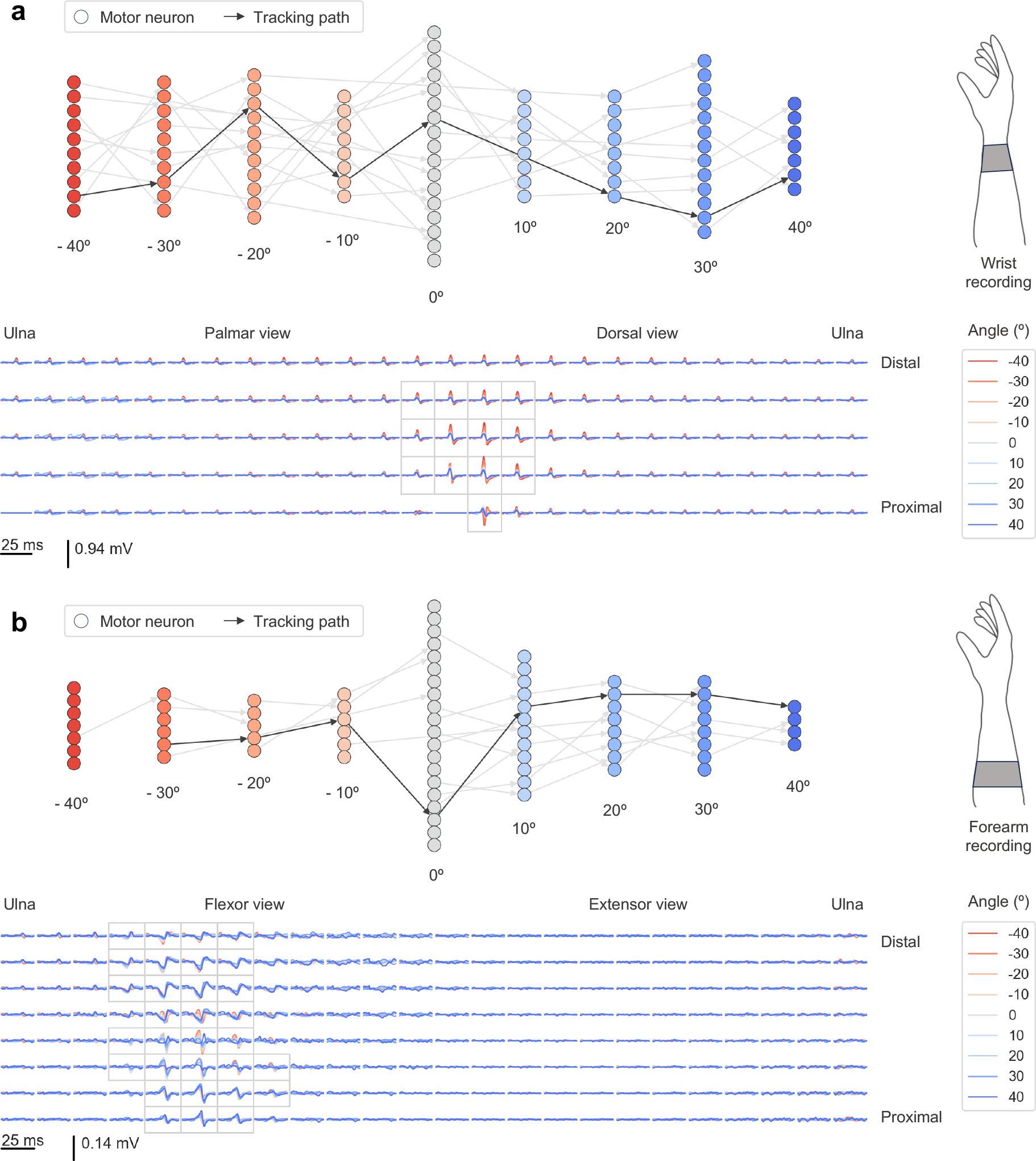
Tracked experimental motor unit action potentials (MUAPs) across angles from the wrist (a) and forearm (b). Each panel includes the detected motor neurons (MNs, circles) at each angle (colour coded in red for wrist flexion, and blue for wrist extension) for a representative finger press and subject. The grey arrows indicate the tracking paths between MUAPs across angles, with the highlighted one (in black) corresponding to the MUAPs display below for the wrist and forearm, respectively. The grey frames indicate the channels that were used for the MUAP comparison based on the normalised mean squared error. In both cases, MUAP differences across angles can be appreciated in the non-propagating components of the wrist MUAPs, and around the innervation point of the forearm ones driven by the change in muscle morphology during dynamic contractions.

To validate the decomposition adaptation results with experimental data, MNs were tracked across the constant phases of the staircase wrist joint angle pattern based on the detected MUAPs after calibrating a decomposition model at each phase angle (Figure 3). In general, more MNs were detected at the forearm than at the wrist (13 (9-19) MNs vs 23 (8-34) MNs, respectively, *p* = 0.004) per subject and finger press after merging MNs across trials and across flexion and extension contractions (Figure 8.a). MNs at the forearm exhibited lower firing rates than those at the wrist (8.94 (6.16-10.55) pps vs 13.06 (10.84-15.30) pps, respectively, *p <* 0.001, Figure 8.b). Decomposition quality was higher at the forearm than at the wrist in terms of PNR (43.25 (40.99-45.79) dB vs 39.62 (37.38-42.35) dB, respectively, *p <* 0.001, Figure 8.c), although the overall CoV was also higher at the forearm than at the wrist (20.30 (15.14-24.58) % vs 19.28 (13.97-24.58) %, respectively, *p* = 0.003, Figure 8.d).

**Figure 8:**
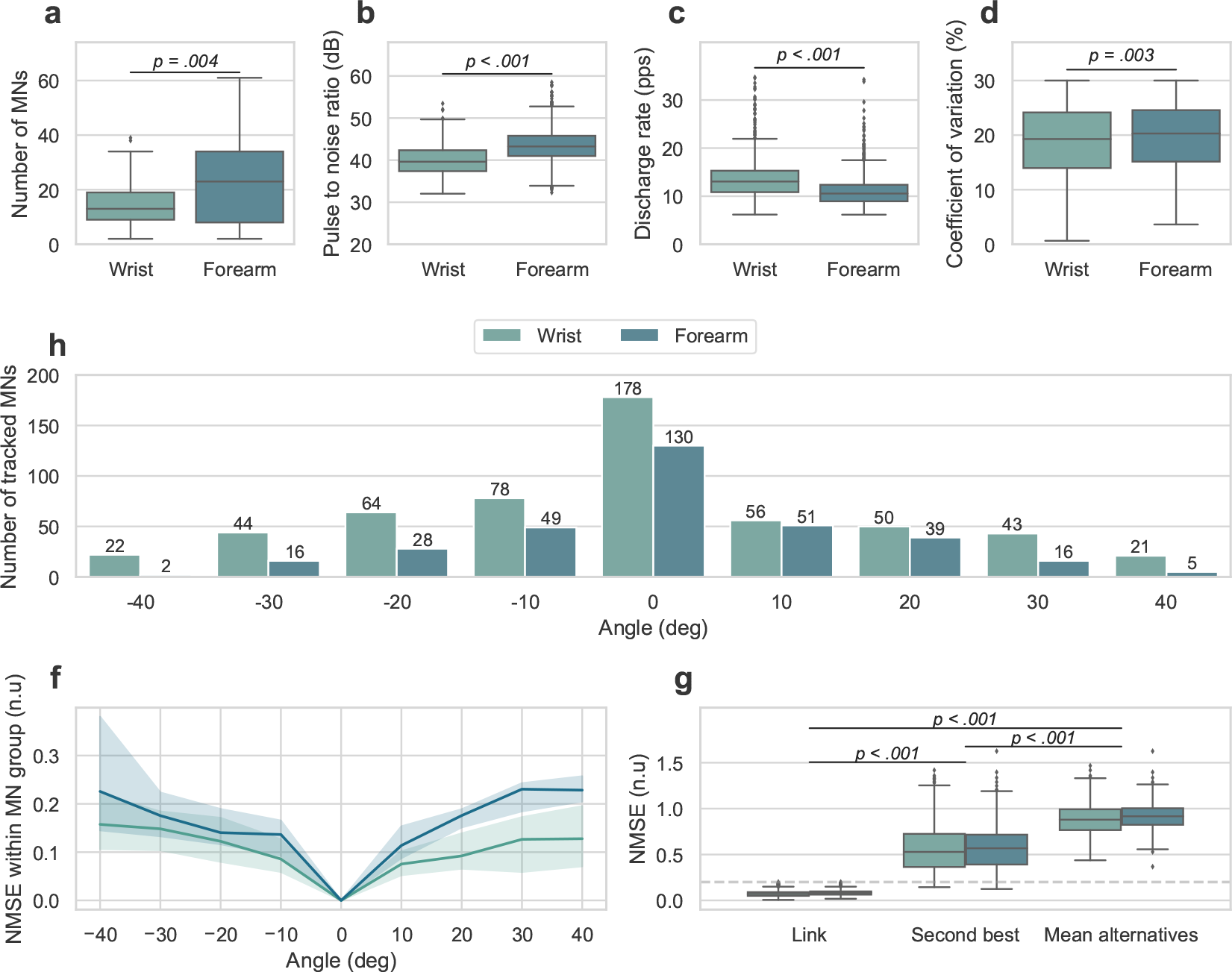
Decomposition calibration for motor unit action potential (MUAP) tracking across angles and properties. (a-d) Properties of the experimental motor neuron (MN) dataset used for MUAP tracking after merging repetitions (see Figure 3). Median and quartile distributions of the number of reliably detected motor neuron (MNs) for each subject across fingers, trials and angles for the wrist (green) and forearm (blue) (a), along with their corresponding pulse to noise ratio (PNR) (b), discharge rate (DR) (c), and coefficient of variation (CoV) (d). Significant *p*-values between the properties at the forearm and the wrist are shown in (a-d) as assessed by the Mann-Whitney U test. After applying the grid-search tracking algorithm, only the MNs that appeared at the reference position (0 °) and at least another angle were kept for further analyses. (h) Depicts the resulting number of tracked MNs for the wrist (green) and forearm (blue) (values at 0°) and the number of MNs present at each angle. This constitutes the dataset for decomposition adaptation validation. (f) Normalised mean squared error (NMSE) between the tracked MUAPs within MN group with respect to the one at 0° for the wrist (green) and forearm (blue), showing an increase in dissimilarity with the change in joint angle. (g) NMSE values between the links across MUAPs at different angles, the second closest MUAP per comparison, and the mean across all possible candidates per comparison for the wrist (green) and foreram (blue). The dashed grey line indicates the assignment threshold for the grid-search tracking algorithm. Significant *p*-values between the three link types are indicated in the plot as assessed by a generalised linear model with repeated measures with location as between-group factor and link types as within-group factor with subsequent pairwise comparisons with Bonferroni correction. The linked MUAPs across angles were significantly more similar (lower NMSE, *p <* 0.001) than the rest of the potential candidates supporting their assignment to the same MN group.

This population of MNs were tracked across angles using the proposed grid-search algorithm based on their MUAPs NMSE. Only the MNs that were tracked for at least two angles (one being 0°as it was the calibration angle for the decomposition adaptation) were kept for further analyses. In total, 178 MNs met these conditions at the wrist and 130 MNs met them at the forearm. Figure 8.h shows the overall number of MNs present at each angle for the wrist and forearm. While more MNs were initially detected at the forearm, MNs at the wrist were tracked over more angles overall (3 (2-3) tracked angles vs 3 (2-4) tracked angles for the forearm and wrist, respectively, *p <* 0.001). This can be explained by the lower MUAP NMSE across angles with respect to 0° at the wrist than at the forearm (Figure 8.f). While the grid-search algorithm did not have a validation set, the NMSE values of the linked MUAPs were significantly lower than those of the second best possible candidates (0.07 (0.05-0.09) n.u vs 0.54 (0.37-0.72) n.u., respectively, *p <* 0.001) and the mean of all possible candidates (0.07 (0.05-0.09) n.u vs 0.89 (0.78-1.00), respectively, *p <* 0.001), showcasing that the selected tracking paths were outliers in the MNs’ (dis)similarity distributions. Furthermore, no significant differences were found between the tracking NMSE between the forearm and the wrist (*p >* 0.05).

### 3.3 Decomposition adaptation in experimental data

The tracked MN firings of the previous section (see Figure 8.h) were used to validate the performance of the decomposition adaptation in experimental contractions. Figure 9.a shows the wrist joint angle for a representative contraction with the corresponding individual finger forces. Figure 9.b depicts the smoothed discharge rates of two of the detected MNs before (grey) and after (blue) adaptation at the wrist along with the tracked ones (red). The detected spike trains over the shadowed part of the contraction in Figure 9.a-b are shown in Figure 9.c for the tracked, adapted, and not adapted decompositions. The figure shows a decrease in decomposition accuracy during dynamic contractions (particularly after 20° in this case) and how decomposition adaptation increased the RoA with the tracked ones.

**Figure 9:**
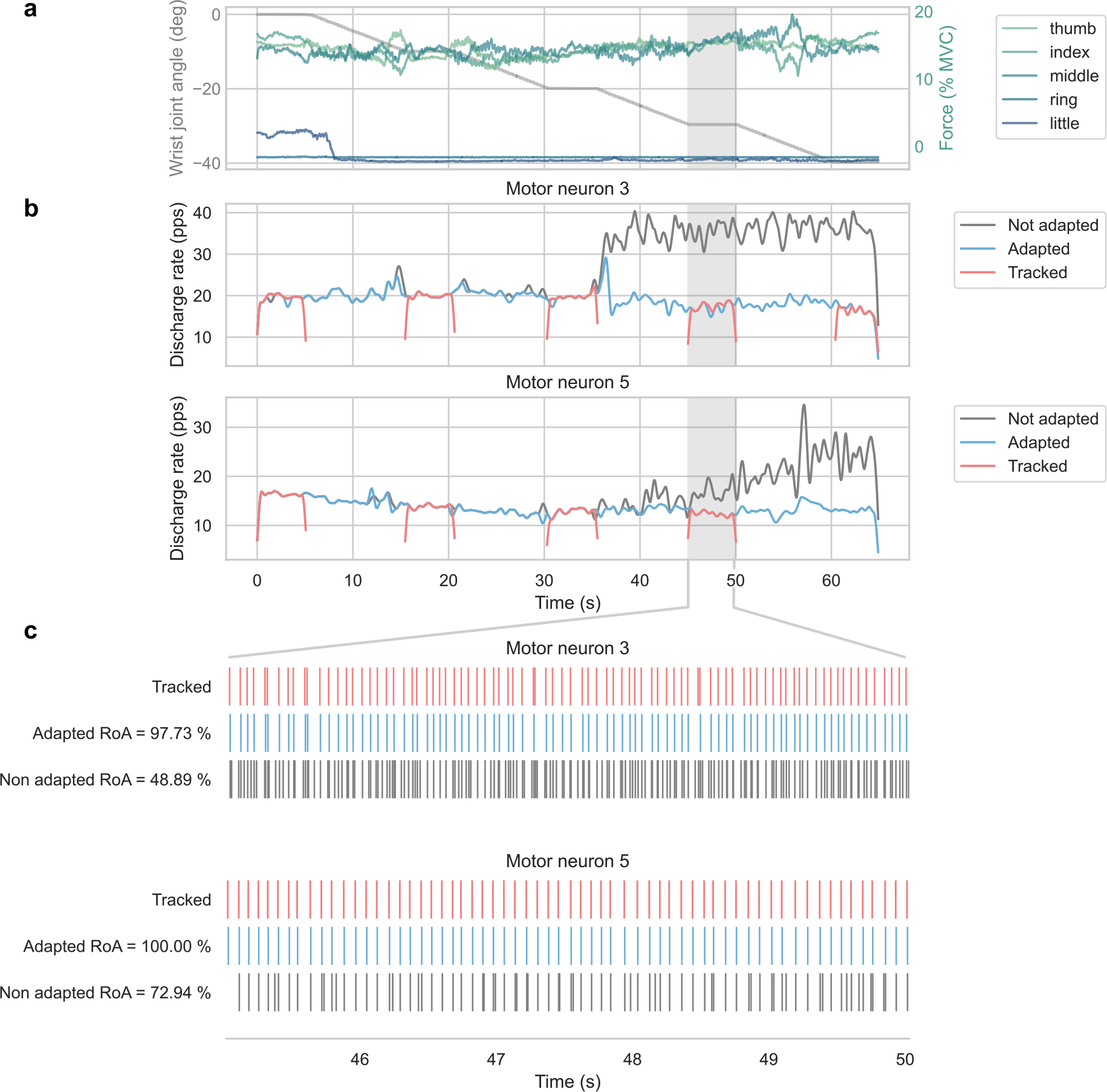
Detection of motor neuron (MN) firings during experimental dynamic contractions from the wrist. (a) Recorded wrist joint angle from the motor encoder (grey) and individual finger forces (colour) for a representative contraction. (b) Smoothed discharge rate of two MN before (grey) and after adaptation (blue) detected at the wrist, along with the smoothed discharge rate of the correspondingly tracked MN based on their MUAP waveforms (red). (c) Spike trains of the tracked MNs (red), and detected firings before (grey) and after adaptation (blue) of the MNs in (b) over the shaded part of the contraction in (a-b). The respective rates of agreement (RoA) for each unit and decomposition approach are shown on the left. The figure illustrates the decrease in decomposition accuracy during dynamic contractions, in this case beyond ∼ 20°, and how decomposition adaptation can restore it.

The overall RoA, sensitivity, precision, NMSE, and PNR for all experimental MNs across angles are displayed in Figure 10 for the wrist (left) and forearm (middle). The median and quartile distributions of the average performance metrics per MNs for both locations before and after adaptation are depicted on the right. Generalised linear models with repeated measures were used to assess the effect of adaptation (within-group factor) and location (between-groups factor), with subsequent pairwise comparisons with Bonferroni correction. Decomposition adaptation significantly improved all performance metrics (*p <* 0.001), with no significant difference between locations (*p >* 0.05). Overall decomposition adaptation increased the median RoA by 15.05% at the forearm (83.94 (63.21-97.75) % vs 98.99 (97.22-100.00) % before and after adaptation respectively, *p <* 0.001) and by 10.67% at the wrist (88.01 (66.26-97.15) % vs 98.68 (95.82-99.40) % before and after adaptation respectively, *p <* 0.001). While an initial statistically significant interaction was found between adaptation and location (*p* = 0.024) no statistically significant differences were found between the effects’ pairwise comparisons after Bonferroni correction (*p >* 0.05 for all cases). A similar effect was observed for the sensitivity, for which decomposition adaptation increased the median sensitivity by 11.28 % at the forearm (88.18 (64.43-98.87) vs 99.46 (98.59-100) % before and after adaptation, respectively, *p <* 0.001) and by 4.98 % at the wrist (94.29 (73.23-98.69) % vs 99.27 (98.32-100.00) % before and after adaptation, respectively, *p <* 0.001). While the interaction between adaptation and location was initially significant (*p* = 0.003), no statistically significant differences were found between the effects’ pairwise comparisons after Bonferroni correction (*p >* 0.05 for all cases).

**Figure 10:**
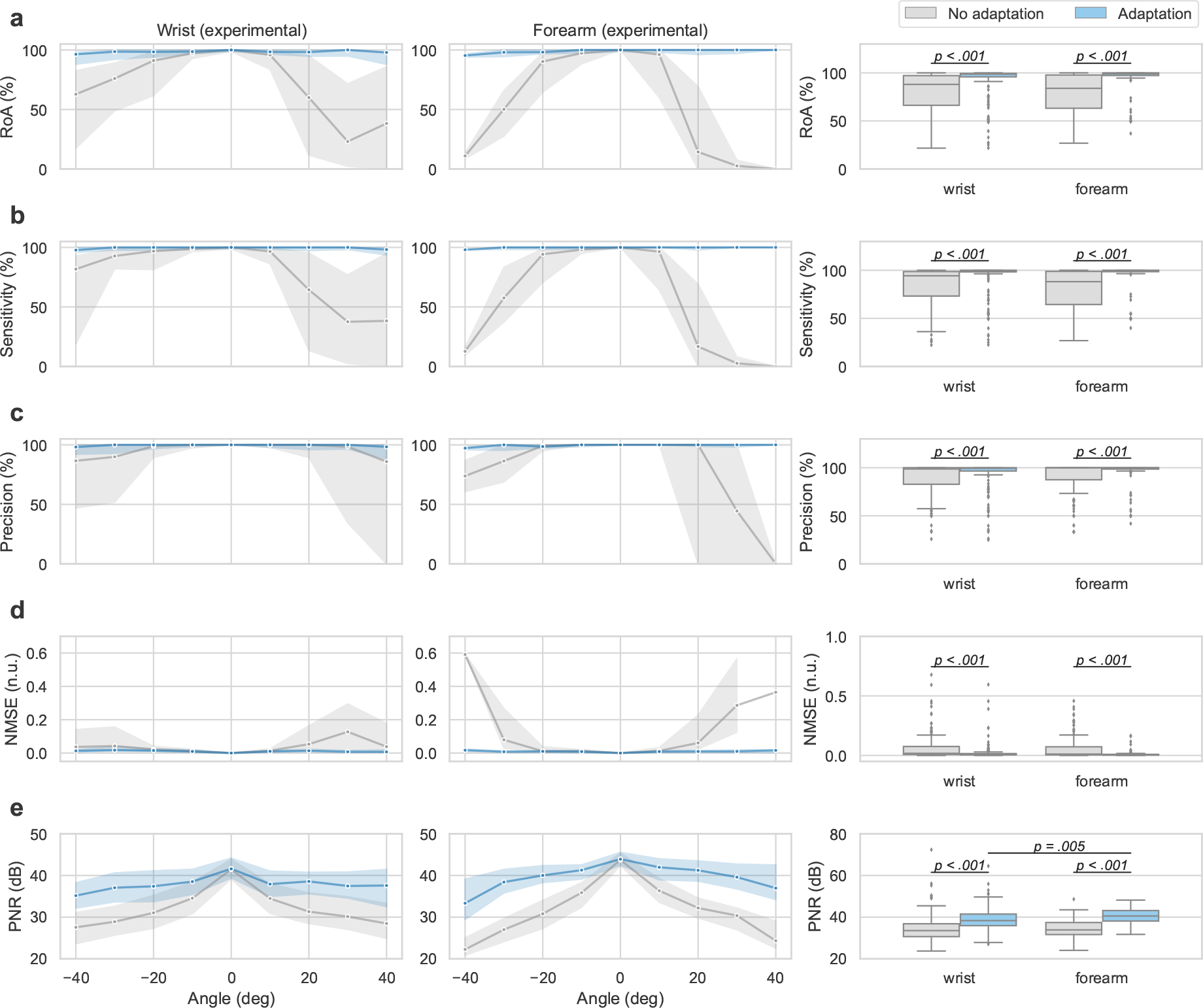
Decomposition adaptation performance in experimental data across the tracked motor neurons (MN). First two columns: median (solid line) and interquartile range (shaded region) before (grey) and after (blue) decomposition adaptation across all tracked MN for each angle for (a) rate of agreement (RoA), (b) sensitivity, (c) precision, (d) normalised mean squared error (NMSE), and (e) pulse to noise ratio (PNR) at the forearm and wrist, respectively. The right panels show the median and quartiles distribution of the metrics across all the tracked MN at the wrist and forearm, respectively. Note the different scale on the y-axis between the line and boxplots for visualisation purposes. Significant *p*-values are show in the plots as assessed by generalised linear models with repeated measures to assess the impact of adaptation (within-group effect) and location (between-groups effect), with subsequent pairwise comparisons with Bonferroni correction. Decomposition adaptation significantly improved all performance metrics (*p <* 0.001), with no significant difference between locations (*p >* 0.05).

**Figure 11:**
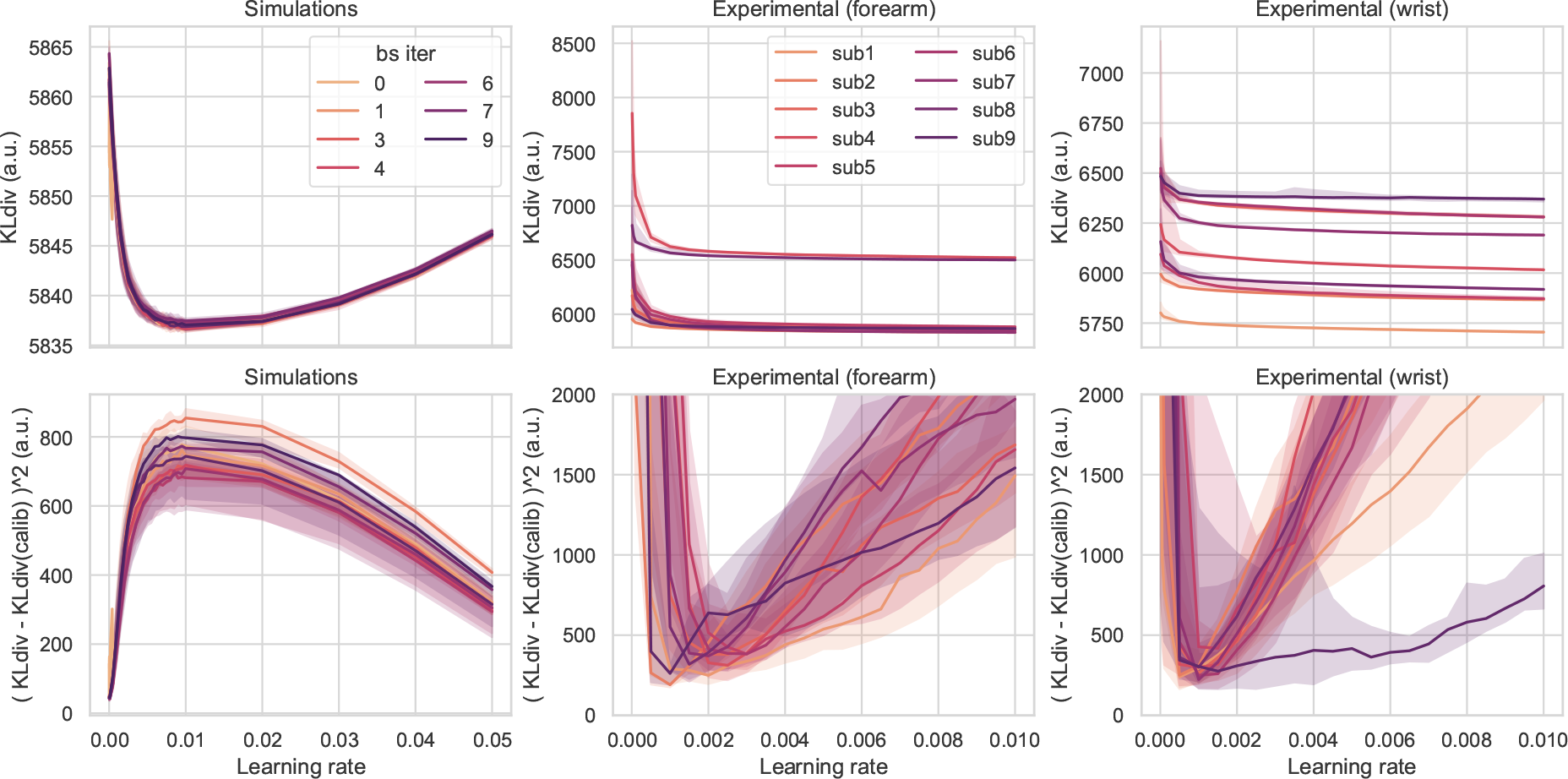
Whitening learning rate optimisation for simulated and experimental data. Top row: median (solid line) and interquartile range (shaded area) of the Kullback-Liebler divergence as a function of the learning rate for each bootstrapping iteration (colour) of the simulated data (left), each subject (colour) at the forearm (middle) and wrist (right). Bottom row: Respective median (solid line) and interquartile range (shaded area) of the median squared error of the Kullback-Liebler divergence between the operational and calibration conditions as a function of the learning rate. While in simulations a global minimum was found for the divergence, the experimental data required the minimisation of the median squared error. Since the error was assumed constant across the contraction, both approaches share the same adaptation rule, albeit a different hyperparameter selection criteria.

Overall, decomposition with fixed parameters from the calibration maintained a high median precision at both forearm (100 (87.50-100.00) %) and wrist (98.73 (82.82-100) %). Still, significant differences in the distribution were found before and after adaptation at both the forearm (100 (98.60-100) %, *p <* 0.001) and the wrist (99.48 (96.57-100) %, *p <* 0.001). In this case, no statistically significant interaction was found between adaptation and location (*p >* 0.05). A similar trend was observed for the NMSE where a significant difference between the distributions was found both at the forearm (0.01 (0.01-0.07) n.u. vs 0.01 (0.00-0.01) n.u. before and after adaptation, respectively, *p <* 0.001) and at the wrist (0.02 (0.01-0.08) n.u. vs 0.01 (0.01-0.02) n.u. before and after adaptation, respectively, *p <* 0.001). As for precision, no statistically significant interaction was found between adaptation and location (*p >* 0.05). Finally, decomposition adaptation increased the median PNR by 6.67 dB at the forearm (33.88 (31.65-37.43) dB vs 40.55 (38.13-43.17) dB before and after adaptation, respectively, *p <* 0.001) and by 4.79 dB at the wrist (33.54 (30.67-36.83) dB vs 38.33 (35.92-41.48) dB before and after adaptation respectively, *p <* 0.001). A statistically significant interaction was found between adaptation and location (*p* = 0.002), which led to a significant difference in the distributions of the forearm and the wrist after adaptation (*p* = 0.005, pairwise comparison with Bonferroni correction).

Taken together, these results show that the proposed decomposition adaptation algorithm significantly improved the decomposition performance in experimental data across all metrics, regardless of the location. Importantly, the convergence of the MUAP tracking and decomposition adaptation algorithms, which are independent from each other, constitute a type of two-source validation.

## 4 Discussion

Identifying MN activities under non-stationary conditions from EMG is essential to complete the vision of wearable non-invasive neural interfaces. Here, we have presented a decomposition adaptation algorithm that leverages the Kullback-Liebler divergence and kurtosis of the signals as metrics for online learning, providing a theoretical framework to optimise the algorithm’s hyperparameters and adapt to non-stationarities by minimising the median squared error with respect to the calibration values. The proposed adaptation algorithm was able to compensate the effect of dynamic contractions resulting in ≥ 90% RoA, sensitivity, and precision (median *>* 98%) in ≥ 80% of the cases in both simulations and experimental data after MN tracking in a two-source validation approach. Below we discuss the relevance of these results for MN detection and interfacing applications over dynamic contractions.

### 4.1 Effect of dynamics in the MUAPs

When muscle fibres shorten and lengthen, their morphology changes, deforming their surrounding tissue, and resulting in varying potentials that dynamically mix the underlying MN firings. In this study, we quantified this effect as changes in the MUAP waveforms with the wrist joint angle in both simulated and experimental conditions (Figure 4 and Figure 7, respectively). While, the NMSE increased with the wrist joint angle deviation from the reference position at 0° in both cases, different nonlinear trends were observed between the simulated and experimental data. This discrepancy can be explained by 1) the effect of spike triggered averaging in the detection of the experimental MUAPs, and 2) the activation of more than one muscle in experimental conditions whose rate of change may not necessarily be the same. However, isolating the contributions of individual muscles from surface recordings is challenging due to the presence of cross-talk [40]. Furthermore, while the effect of spike triggered averaging could have been reduced by recording longer contractions at more wrist joint angles, a trade-off was necessary to avoid participants’ fatigue. Importantly, the maximum median NMSE (∼ 0.2 n.u.) was similar in both simulated and recorded MUAPs, supporting the suitability of NeuroMotion [19] as a tool to model dynamic contractions for dynamic decomposition models. In fact, the simulated/detected MUAP changes were qualitatively similar to those reported by Holobar and collaborators [12, 13] for the biceps brachi. However, a direct quantitative comparison is complicated due to the different targeted muscles and metrics used (cross-correlation coefficient vs NMSE).

While the cross-correlation coefficient is arguably a more common metric for MUAP tracking in isometric contractions [41, 42, 43], its use during dynamics should be treated with caution due to its insensitivity to amplitude changes. While Oliveira and Negro [15] pioneered its used to track MUAPs from the tibialis anterior between dynamic ankle dorsiflexions, their success may be explained by the limited range of motion explored (∼ 20°). In fact, a recent study has shown that decomposition performance for the tibialis anterior over dynamic contractions with up to 20° was equivalent to that of isometric contractions in hybrid data [14]. Since in this study we explored a substantially larger range of motion (80°) over smaller muscles at the forearm, here we used the NMSE instead. Notably, NMSE can reflect amplitude changes and was previously proposed as a metric to detect the unique representation of MUAPs in surface recordings [39]. This sensitivity was important to 1) characterise the overall MUAP changes during the simulated/experimental contractions to provide benchmarks for the dynamic modulation compensated by the decomposition adaptation, and 2) to ensure that the uniqueness of different MUAPs simulated by NeuroMotion did not collapse when inducing the dynamic modulation (Figure 4.a-g). Therefore, when coupling the NMSE with the proposed channel selection approach that identifies the channels with outlying peak amplitude, the resulting metric condenses temporal, spatial, and amplitude features to measure (dis)similarities. It is worth noting that channel selection is essential over high-density recordings to avoid deceivingly large similarities due to a large amount of common silent channels as those present in Figure 7.

Experimental MUAP tracking showed that the modulation induced by muscle fibre shortening/lengthening was higher at the forearm than at the wrist as assessed by the NMSE across angles. This is attributable to the convergence of most muscle fibres into tendons at the wrist, who exhibit different contractile properties. Importantly, the MUAPs at the wrist are dominated by non-propagating far-field potentials (Figure 7.a) that emerge during the progressive extinction of the action potential at the muscle-tendon interface [8, 9, 10]. Therefore, these MUAPs are arguably more susceptible to relative displacements of the muscle-tendon interface than to the contractile properties of the muscle. Conversely, at the forearm, these changes alter the propagating MUAP components and relative location of the innervation zone (Figure 7.b) resulting in larger NMSE values with respect to the reference position. The higher sensitivity of recordings over the muscle belly to dynamic contractions may explain why although more MNs were decomposed at the forearm than at the wrist, the latter ones were tracked along more angles. Yet, these results rely on the performance of the tracking algorithm. While the tracking approach did not have a validation set per se, the linked MUAPs exhibited a NMSE significantly lower than that of their corresponding second best alternative, supporting the hypothesis that the representation of MUAPs belonging to the same MNs are more similar within themselves than across different MNs. It is also worth noting that by sequentially linking the most similar MUAPs across angles, the tracking algorithm only assumes local continuity in the (dis)similarity distribution and can potentially track larger changes than the predefined threshold with respect to the reference position (in this case at 0°, see Figure 8.f). Here we set the NMSE linking threshold to 0.2 n.u. as an equivalent dissimilarity value of the correlation coefficient threshold previously used in the literature (0.8, [15]). While arguably the selection of this threshold may condition the results, with potential for further improvement, the fact that decomposition adaptation resulted in ≥ 90% RoA, sensitivity, and precision with respect to the firings of the experimentally tracked MUAPs in ≥ 80% of the cases, supports the accuracy of the grid search tracking approach in a form of two-source validation since both methodologies are completely independent from each other.

However, it should be noted that the lower number of MNs tracked at each angle with respect to the total ones detected at 0° does not necessarily reflect a decreased recruitment of MNs during dynamics. In fact, the findings of Oliveira and Negro [15] suggest that MN rate coding (as oppose to MN recruitment) is the main driver of muscle shortening and lengthening during small dynamic ankle dorsiflexions at different speeds and constant load. Instead, the lower number of MNs tracked at each angle echo the fact that decomposition algorithms tend to identify only a portion of the active MNs. Moreover, here we only used the reliably detected MNs for tracking with PNR ≥ 30 dB, 5 ≤ DR ≤ 35 pps, and CoV ≤ 30 %. The selected PNR threshold has been shown to be equivalent to a *>*90% sensitivity and *<*2% false alarm rate in the decomposition accuracy [33]. Similarly, the aforementioned DR and CoV ranges correspond to the physiological behaviour of MN firings [24, 37] and their accurate identification [38]. However, the imposition of such constrains likely decreased the overall number of detected MNs that may otherwise have been kept if manual validation was applied [44], reducing our ability to track them across angles. Nonetheless, manual decomposition editing is not possible in human-machine interaction applications, and the overall number MNs selected for the validation of the decomposition adaptation (308 MNs in total, 178 at the wrist, and 130 at the forearm) was higher to the one reported in previous studies with hybrid data sets (252 MNs in [12, 13]) and experimental data (18 MNs in [18]).

### 4.2 Effect of dynamics in the decomposition model

The larger modulation at the forearm during dynamics was also independently observed as a higher median squared error of the Kullback-Liebler divergence between operational and calibration conditions at the forearm than at the wrist (see Supplementary Figure 12). As expected, larger MUAP differences with respect to the calibration conditions affected the signals’ covariance, increasing the divergence error. In fact, such discrepancies resulted in an optimal learning rate twice as high during the whitening adaptation (*η*_2_ = 2*e*^−3^ vs *η*_2_ = 1*e*^−3^ for the forearm and wrist, respectively). Notably, once the whitening adaptation was applied to compensate the effect of the dynamics, the same optimal learning rate for the separation vectors was found in both locations (*λ* = −5*e*^−4^). This supports the efficacy of the proposed adaptation algorithm, since the minimisation of the median squared error of the sources’ kurtosis with respect to the calibration relates to the sources’ higher order statistics rather than the dynamics of the volume conductor.

**Figure 12:**
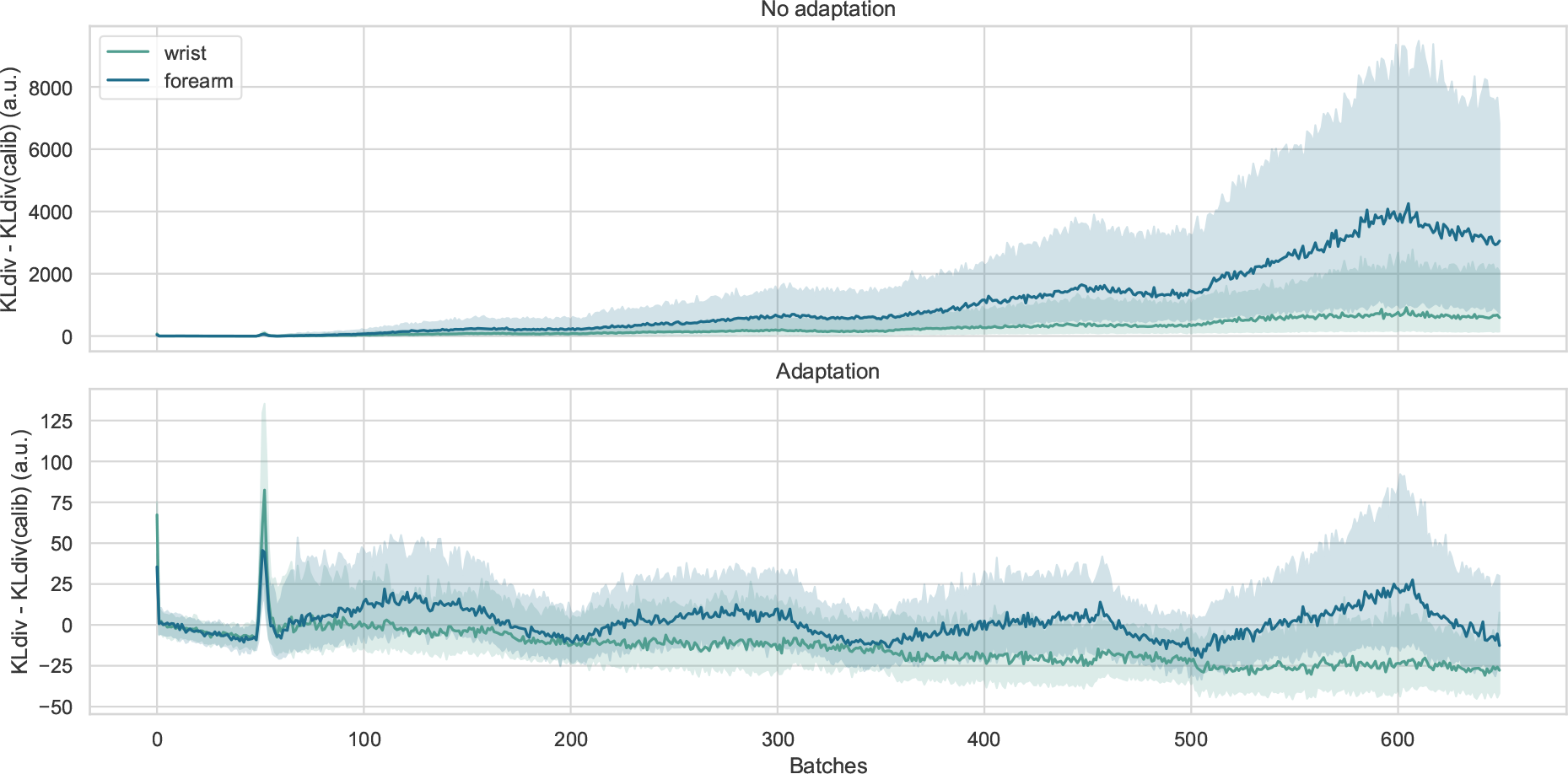
Kullback-Liebler divergence error. Top: Median (solid line) and interquartile range (shaded area) of the Kullback-Liebler divergence error with respect to the calibration before adaptation across all experimental contractions for the wrist (green) and forearm (blue). Bottom: Same as at the top, but after applying the proposed whitening adaptation. Note the change in scale on the y-axis for visualisation purposes. The two peaks at the beginning of the plot correspond to the effect of the decomposition extension factor and an artifact produced by the motor at the initialisation of the motion, respectively.

Still, few considerations need to be taken into account. First, the optimisation of the whitening hyperparameters was slightly different in simulations than in experimental data. The Kullback-Liebler divergence was minimised in simulated conditions as opposed to the minimisation of the median squared error with respect to the calibration values applied to the experimental data. This discrepancy can be explained by the different optimisation topology between the simulated and the experimental data (Supplementary Figure 11). These differences could be attributable to the activation of a single muscle in simulations, the injection of artificial noise, or the dynamics of the volume conductor model, which are all factors that can alter the covariance matrix. However, it is worth noting that here we assumed a constant error rate during both the whitening and separation vector optimisation, i.e. the optimised parameters correspond to the product of the learning rates and the error, either the divergence or kurtosis, and therefore the adaptation rules for both the simulated and experimental data are the same, except for the choice of the whitening hyperparameter. This was a simplification assumption, yet the inclusion of the actual error term in the adaptation may provide further benefits such as faster learning or the ability to tune each MN separation vector differently due to varying error rates in the kurtosis of the sources within a given contraction. However, the consistency of the minimisation of the median squared error across subjects and locations was remarkable (Supplementary Figure 11), supporting the validity of such assumption for the current task.

### 4.3 MN identification performance during dynamics

Decomposition performance results showed that without adaptation, the median RoA starts decreasing after 10°deviations in both extension and flexion for simulations and experimental data. To the best of our knowledge, this is the first study to characterise the effect of muscle shortening and lengthening (here determined as the wrist joint angle during flexion and extension) in the performance of the decomposition (without adaptation) for both simulated and experimental data across a wide range of motion (80°). While some MN firings were accurately detected over larger deviations during wrist flexion, from 30° performance was lost overall, particularly at the forearm. This is consistent with the larger MUAP changes observed at the forearm than at the wrist with the angle deviation. The found tolerance of the decomposition model to changes in posture is akin to the one observed in previous studies for the tibialis anterior [15, 14] and MUAP similarity in the biceps brachii [13]. Here we included angle plateau phases in the dynamic contraction to enable the generation of ground truth via MUAP tracking from independently calibrated decomposition models at each plateau angle. However, these results supports the validation of future decomposition adaptation approaches over purely dynamic contractions as long as the decomposition model is calibrated over segments within the decomposition postural tolerance (∼10°).

The proposed adaptation significantly improved all performance metrics (RoA, sensitivity, precision, NMSE, and PNR), resulting in ≥ 90% RoA, sensitivity, and precision in ≥ 80% of the cases in both simulations and experimental data (median *>*98% for the three metrics across data sets). Importantly, no statistically significant differences between the wrist and the forearm were found for all metrics, showcasing the ability of the algorithm to compensate different modulations related to the anatomical factors described before. In addition, the performance of the proposed decomposition adaptation was equivalent if not higher than previously proposed adaptation approaches based on recursive updates of the decomposition parameters. For instance, the approach proposed by Chen et al. [17] resulted in average ∼ 80% sensitivity and precision when compensating MUAP modulations at different speeds simulated in a cylindrical volume conductor. More recently, Yeung et al. [18] reported an average ∼ 89% RoA in experimental data over a moderate range of motion (up to 25% wrist extension, equivalent to ∼17° based on the ∼ 70° maximum range of motion of the wrist during extension [30]) after adaptation. While direct comparison is difficult due to the different conditions analysed, the achieved performance over the extensive range of motion evaluated here (−40° flexion to 40° extension, corresponding to the range needed for activities of the daily living [29]) arguably represents more challenging conditions for the decomposition. Importantly, the aforementioned approaches included ad-hoc values for the learning rates and recursive support, which are conceivably hard to transfer and tune to different data sets without ground truth. Here instead, we proposed two online learning metrics based on the Kullback-Liebler divergence and kurtosis of the signals to overcome this limitation.

### 4.4 Translational potential, limitations, and future directions

The overarching motivation of the proposed decomposition adaptation was to reliably detect previously identified MUs (in this case, under isometric contractions) during non-stationary conditions such as postural changes or dynamic contractions. The ultimate goal was to develop an adaptation method that would be suitable for human-machine interaction applications with the potential to improve the overall robustness of the neural interface. Importantly, the enhanced stability of the MU identification could also result in benefits during motor control by promoting learning [45]. To ensure this compatibility, the proposed decomposition adaptation had not only to be accurate but also execute under the neuromechanical delay (∼ 200 ms, [23]). In this case, once the optimal learning rates were identified, the total execution time of the entire adaptation pipeline was ∼22 ms per 100 ms batches (approximately half of the previously proposed approaches [17, 18]), showcasing the suitability of the proposed adaptation for real-time human-machine interaction applications.

Crucially, the proposed approach provides two online learning metrics based on the Kullback-Liebler divergence and kurtosis of the signals to tune the algorithm’s hyperparametes. Importantly, these metrics are agnostic to the ground truth and their optimisation in this study has resulted in ≥ 90% RoA, sensitivity, and precision in ≥ 80% of the cases in both simulations and experimental data, demonstrating the ability of these metrics to capture the modulation induced by the dynamic contractions. The success of these metrics is rooted in the assumptions of the blind source separation model that first uncorrelates the mixed signals and subsequently finds the orthogonal transformation that guarantees their independence [32] (or sparsity in the decomposition model [2]). Therefore, the proposed metrics can be considered proxies of decomposition performance where a high Kullback-Liebler divergence indicates lower uncorrelation of the EMG signals; and low kurtosis (estimated here by the logcosh contrast function) reflects a poorer separation of the underlying sources similar to the PNR [33]. While the evaluated decomposition performance was carried out after hyperparameter optimisation, which is arguably a form of model precalibration that requires initial data, the fact that the obtained hyperparameters generalised across subjects and locations shows promising potential for its application to different tasks where ground truth is not available. However, in this study we only evaluated slow staircase contraction patters for wrist flexion and extension, and so the interplay between online learning and dynamic modulation speed still needs to be assessed. Nonetheless, these results open the door to a two double calibration scenario where the decomposition model is initialised first during an isometric contraction, and the underlying dynamic modulation is learnt later in a second tuning phase. Future work should address which approach is more effective and robust for human-machine interfacing.

## 5 Conclusions

Identifying MN activity reliably from EMG under non-stationary conditions is essential to develop robust wearable non-invasive neural interfaces. Here, we introduced a novel decomposition adaptation algorithm that leverages the Kullback-Liebler divergence and kurtosis of the signals as metrics for online learning. The proposed approach provides a ground-truth agnostic theoretical framework to tune the algorithm’s hyperparameters and adapt to non-stationarities, such as dynamic contractions, by minimising the median squared error of the metrics with respect to the calibration values in real-time (∼ 22 ms execution time per 100 ms batches). The proposed adaptation algorithm significantly improved all decomposition performance metrics with respect to no adaptation in a wide range of motion of the wrist during flexion and extension (80°). Results showed ≥ 90% RoA, sensitivity, and precision in ≥ 80% of the cases in both simulations and experimental data from the wrist and forearm after tracking MNs across wrist joint angles based on their MUAP waveforms in a two-source validation approach. These findings show promising potential for the application of the proposed decomposition adaptation approach to different tasks where ground truth is not available, and real-time human-machine interfacing.

## Acknowledgements

This work was supported by the Engineering and Physical Sciences Research Council—Centre for Doctoral Training in Neurotechnology for Life and Health (IMG), Meta (IMG), the European Research Council, Synergy Grant—project Natural BionicS, No. 810346 (DYB, DF), and the Imperial-META Wearable Neural Interfaces Research Centre (DYB, DF). This study utilised the expertise and equipment at the Imperial College Advanced Hackspace.

## Ethical statement

This experimental protocol was approved by the local ethics committee of Imperial College London (reference number 18IC4685) and was conducted in accordance with the Declaration of Helsinki. All participants signed informed consent forms before the experiment.

## Data availability

The data that support the findings of this study are available upon reasonable request from the authors.

## Supplementary figures

